# Single-cell Long Non-coding RNA Landscape of T Cells in Human Cancer Immunity

**DOI:** 10.1101/2020.07.22.215855

**Authors:** Haitao Luo, Dechao Bu, Lijuan Shao, Yang Li, Liang Sun, Ce Wang, Jing Wang, Wei Yang, Xiaofei Yang, Jun Dong, Yi Zhao, Furong Li

## Abstract

The development of new therapeutic targets for cancer immunotherapies and the development of new biomarkers require deep understanding of T cells. To date, the complete landscape and systematic characterization of long noncoding RNAs (lncRNAs) in T cells in cancer immunity are lacking. Here, by systematically analyzing full-length single-cell RNA sequencing (scRNA-seq) data of more than 20,000 T cell libraries across three cancer types, we provide the first comprehensive catalog and the functional repertoires of lncRNAs in human T cells. Specifically, we developed a custom pipeline for *de novo* transcriptome assembly obtaining 9,433 novel lncRNA genes that increased the number of current human lncRNA catalog by 16% and nearly doubled the number of lncRNAs expressed in T cells. We found that a portion of expressed genes in single T cells were lncRNAs which have been overlooked by the majority of previous studies. Based on metacell maps constructed by MetaCell algorithm that partition scRNA-seq datasets into disjointed and homogenous groups of cells (metacells), 154 signature lncRNAs associated with effector, exhausted, and regulatory T cell states are identified, 84 of which are functionally annotated based on co-expression network, indicating that lncRNAs might broadly participate in regulation of T cell functions. Our findings provide a new point of view and resource for investigating the mechanisms of T cell regulation in cancer immunity as well as for novel cancer-immune biomarker development and cancer immunotherapies.

## Introduction

T cell checkpoint inhibition therapies, such as targeting exhausted CD8^+^ T cells and regulatory T cells (Tregs), have shown remarkable clinical benefit in many cancers [1–3]. Nevertheless, the mechanisms underlying therapy response or resistance are largely unknown, which leads to the different therapeutic efficacies among cancer patients [4–8]. To better understand the mechanisms that underlie successful response to immunotherapy, more comprehensive studies to explore the whole transcriptome of individual T cells in tumor ecosystems are desired. Long non-coding RNAs (lncRNAs), defined as a class of non-coding RNAs longer than 200 nucleotides with no or low protein-coding potential, comprise a large proportion of the mammalian transcriptome [9–12]. Accumulating evidence has suggested that lncRNAs are widely expressed in immune cells and play crucial roles in cancer immunity by regulating the differentiation and function of T cells [13–17]. For example, overexpression of *NKILA*, an *NF*-κ*B*-interacting lncRNA, correlated with T cell apoptosis and shorter patient survival [18], and an enhancer-like lncRNA *NeST* regulates epigenetic marking patterns of *IFN*-γ-encoding chromatin and induce synthesis of *IFN*-γ in CD8 T cells [19]. However, previous studies seem to be somewhat scattered and the landscape and comprehensive functional analysis of lncRNAs in T cells in cancer immunity are still lacking.

The dramatic advances of single-cell RNA sequencing (scRNA-seq) technologies have gained unprecedented insight into the high diversity in T cell types and states compared to bulk RNA sequencing methods, which do not address the complex structures of tumor microenvironment [20–25]. Despite the advantages of single-cell resolution, in current most scRNA-seq studies of cancer immunology have generally focused on coding genes, overlooking the large amounts of lncRNAs. Detailed understanding of lncRNAs at the single-cell level was challenging owing to their relatively low and cell-specific expression [26–28]. As a widely used scRNA-seq approach, 3’-end sequencing technologies such as droplet-based 10X Genomics have lower RNA capture efficiencies, leading to the dropout evets and technological noise for lowly expressed lncRNAs [29]. Furthermore, accurate identification of novel lncRNAs is not suitable for the 3’-end sequencing technologies, but such analysis could be achieved by using full-length scRNA-seq technologies such as SMART-seq2 [30]. In addition, the sampling noise in scRNA-seq is generated through sampling of limited RNA transcripts from each cell [31], leading to a highly noisy estimation for most lncRNAs. Therefore, to effectively characterize the lncRNA landscape at single-cell level, attention should be paid to choosing appropriately scRNA-seq data and analytical approaches.

Here, using unprecedentedly large-scale full-length single-cell transcriptome data of more than 20,000 T cells from various tissues across three cancer types, we created a full annotation of the T cell lncRNA transcriptome and analyzed the functional roles associated with different T cell states. Our study aims to provide a basic and valuable resource for the future exploration of lncRNA regulatory mechanisms in T cells, which may facilitate novel cancer-immune biomarker development.

## Results

### *De novo* transcriptome assembly of lncRNAs from scRNA-seq data of T cells

To investigate the landscape of human lncRNAs in T cells across different tissues, patients and cancer types, we collected the data of 24,068 T cells (the size of the gzip-compressed FASTQ file was 7.5 TB) generated by full-length single-cell RNA sequencing with SMART-seq2, including the raw data of 9,878 cells from colorectal cancer (CRC) patients (2.8 TB), 10,188 cells from non-small-cell lung cancer (NSCLC) patients (3.1 TB), and 4,002 cells from 5 hepatocellular carcinoma (HCC) patients (1.6 TB) [32–34] (Figure S1A and Table S1). These cells were collected from peripheral blood, adjacent normal, and tumor tissue from each patient and sorted into CD3^+^CD8^+^ (CD8) and CD3^+^CD4^+^ (CD4) T cells. The reads of each cell were mapped to the human reference genome (hg38/GRCh38), and the cells with unique mapping rates of less than 20% were removed. The remaining cells with on average 1.04 million uniquely mapped read pairs (0.63 million splices on average) and at least one pair of T cell receptor (TCR) α and β chains enabled us to detect the expressed lncRNAs (Figure S1B-D).

Next, to generate the comprehensive T cell transcriptome beyond the currently reference annotation, we performed *de novo* transcriptome assembly using the StringTie method [35]. Although StringTie could be run by providing the reference annotation to guide the transcript construction, in current study we focused on to what extent it could assemble the whole transcriptome without the prior annotation. Based on the T cell dataset from HCC patients, we first measured the extent of assembly in each T cell and found that an average of 4,752 transcripts could be assembled at single-cell level, and an average of 69.8% (3,318/4,752) were matched to reference models (including reference protein-coding genes from GENCODE v31 and reference lncRNA genes from RefLnc database) (**Figure 1A**).

**Figure 1.**
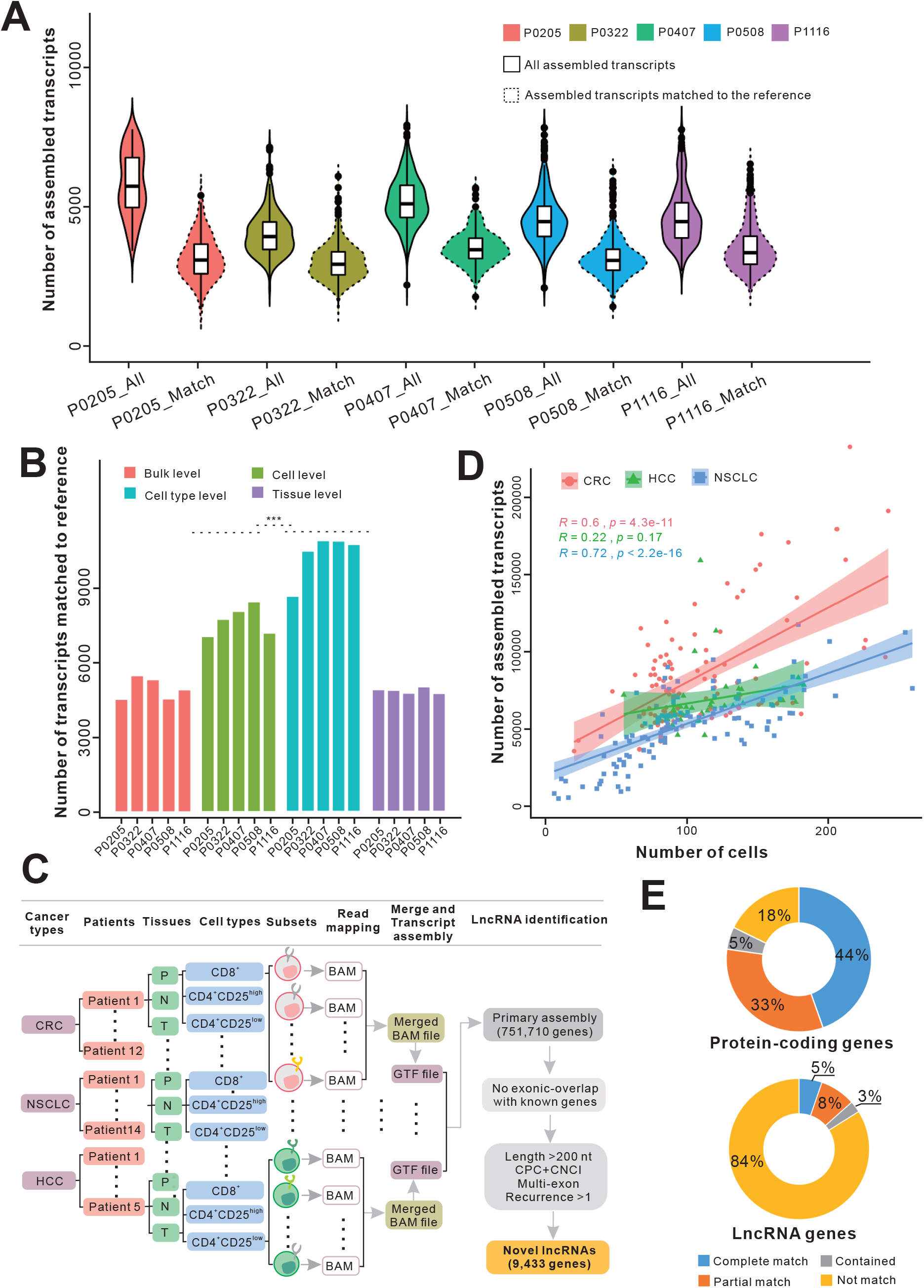
The statistics of assembled transcripts and workflow for novel lncRNA identification process in T cells during cancer immunity. **A**. Violin plots showing the number of assembled transcripts and the number of those matched to the reference at single cell level across five HCC patients. **B**. Number of assembled transcripts that matched to reference across five HCC patients based on four different strategies. *** indicates *P*-value < 0.001 (Wilcoxon rank sum test). **C**. Correlation of the number of cells and the number of assembled transcripts across different subsets for CRC, HCC and NSCLC. A 95% confidence interval was added and shown as coloured regions. **D**. Scheme of pipeline used to identify the novel lncRNAs expressed in T cells during cancer immunity using three full-length scRNA-seq datasets. **E**. The statistics of assembled transcripts that matched to reference protein-coding and reference lncRNA genes. CRC, colorectal cancer; HCC, hepatocellular carcinoma; NSCLC, non-small-cell lung cancer; P, peripheral blood; N, adjacent normal tissue; T, tumor tissue.

To explore the best way to obtain novel transcripts, we compared the assembly results using three different approaches based on HCC dataset: (1) mapping and assembling for each single cell individually (cell-level); (2) assembling transcripts based on merged mapping results from each cell type of each patient (cell type-level); (3) assembling transcripts based on merged mapping results from each tissue of each patient (tissue-level). The transcripts assembled from each approach were merged independently and compared with reference genes respectively (Figure S1E). We found that the number of assembled transcripts matching to reference genes based on the cell type-level strategy (average 105,527 transcripts) was significantly higher than in cell-level or tissue-level methods (average 77,860 and 49,689 transcripts respectively, *P*-value < 0.001, Wilcoxon rank sum test) (Figure 1B). Furthermore, the average number of matched transcripts from the cell type-level was more than twice that from the bulk-seq method (average 48,854 transcripts) (Figure 1B).

According to the cell-type pooling strategy, the cells from all patients across three cancer types were partitioned into 266 subsets (Figure 1C and Figure S1A), and the mapping results of cells from the same subset were merged and fed into assembling program. We found the number of assembled transcripts across different subsets showed positive correlations with the number of cells in these subsets in both CRC and NSCLC datasets (Pearson correlation coefficients = 0.6 and 0.72, *P*-value = 4.3e-11 and < 2.2e-16, respectively), but not in the HCC dataset (Pearson correlation coefficient = 0.22, *P*-value = 0.17) (Figure 1D and Figure S1F). Then, assembled transcripts from all subsets were merged together, and a total of 751,710 primary genes were obtained. Next, we compared our assembled transcriptome with reference gene models. The results showed that reference lncRNAs had a lower detection rate than protein-coding genes. Specifically, 82% (16,399/19,938) of the known protein-coding genes in GENCODE v31 could be verified (44%, 8,893/19,938 were complete match with the same intron chain), while 16% (9,567/59,489) of known lncRNA genes were verified (5%, 3,140/59,489 were complete match) (Figure 1E). These findings suggested that lncRNAs were expressed in a much more cell-specific manner than protein-coding genes and further studies to uncover novel lncRNAs specifically expressed in human T cells were needed.

From the primary assembly, we developed a custom pipeline to identify novel lncRNAs. Briefly, we first selected transcripts that were no shorter than 200 nucleotides and have multiple exons. The transcripts that overlapped with both known protein-coding and known lncRNA genes were filtered out. Then, the transcripts lacking coding potential predicted by both CPC [36] and CNCI [37] utility were retained. Finally, the remaining transcripts that were reconstructed in at least two subsets with complete match were defined as the novel lncRNA catalog (Figure 1C). Through this multi-layered analysis, we identified 9,433 previously unknown lncRNA genes (13,025 transcripts with mean length of 1,112 nucleotides), which increased the number of current human lncRNA catalog [38] by 16% and nearly doubled the number of lncRNAs expressed in human T cells.

Finally, we performed experimental validation to evaluate the robustness of our identified novel lncRNAs. First, fresh peripheral blood samples were collected from three CRC patients (Table S2). Then, mononuclear cells were isolated from each sample. CD8 and CD4 T cells were separated by immunomagnetic beads and the separation efficiency was verified by flow cytometry (**Figure 2A**). Next, we selected 50 novel lncRNAs for quantitative real-time polymerase chain reaction (qRT-PCR) analysis and Sanger sequencing across T cell samples. As a result, 38 novel lncRNAs could be verified successfully by Sanger sequencing (Table S3). As an example, for a novel lncRNA *TCONS_00180551* located in an intergenic region of chromosome 11, the blat search result of Sanger sequencing exactly matches the junction of this novel lncRNA (Figure 2B).

**Figure 2.**
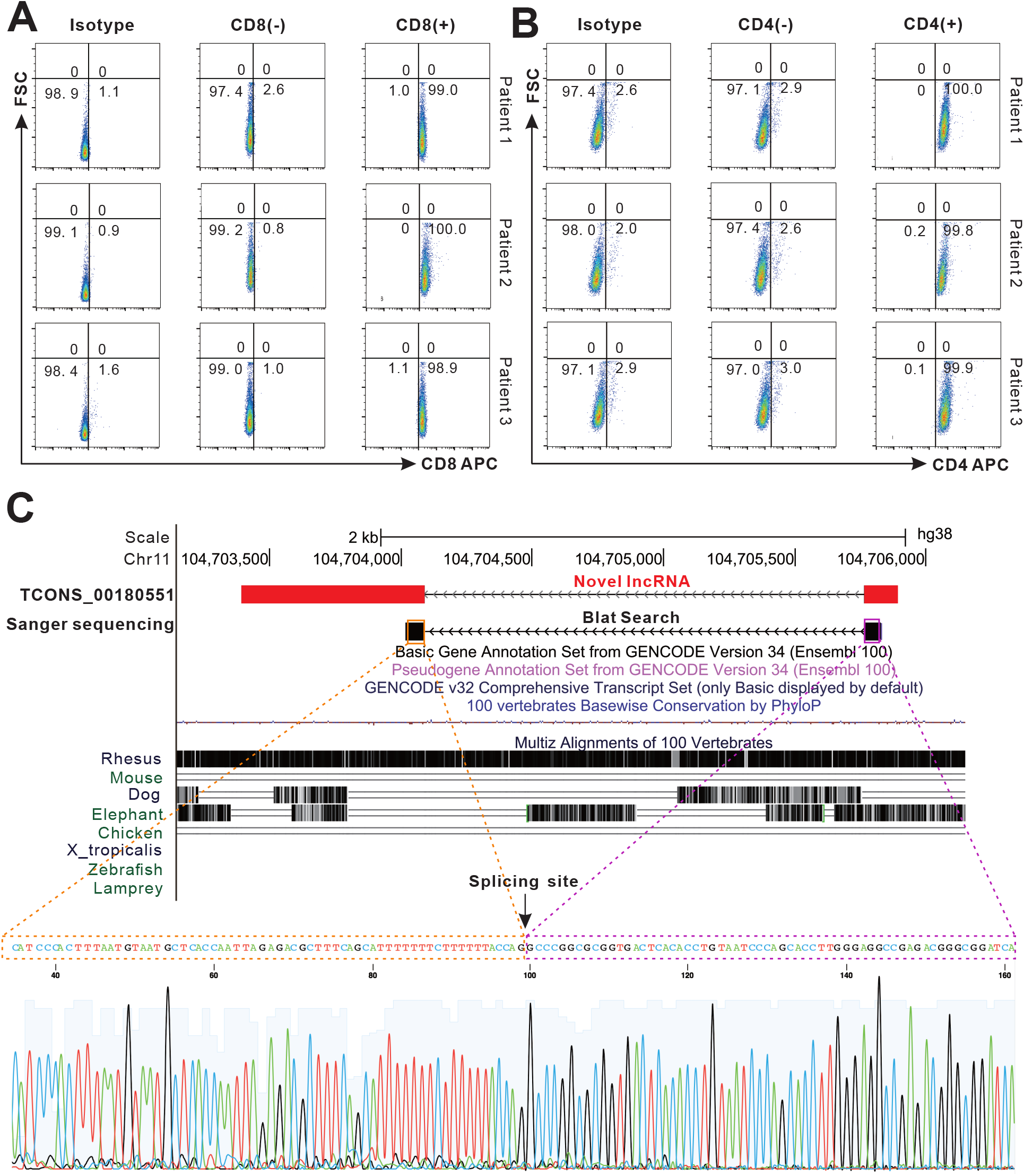
Single T cell sorting and quality evaluation of an example novel lncRNA. **A**. The results of flow cytometric analysis. CD8 and CD4 T cells from three patients were separated by magnetic beads and stained with flow cytometry antibody CD8-APC and CD4-APC respectively (Isotype was used as negative control). **B**. An example of novel intergenic lncRNA that was validated by Sanger sequencing. The genomic views are generated from UCSC genome browser. The spliced sequence outputted by Sanger sequencing is shown.

### The characterization and expression analyses of lncRNAs in T cells

Based on the relative genomic locations to reference protein-coding genes, the novel lncRNAs were classified into three locus biotypes, including 6,525 as intergenic, 3,187 as intronic and 3,313 as antisense lncRNAs. As in the case of reference lncRNAs, these novel lncRNAs showed fewer exons (the average number of exons was 2), lower detection rates and average gene abundance than protein-coding genes at single-cell level (**Figure 3A-B**). Specifically, by using pseudoalignment of scRNA-seq reads to both reference and novel lncRNA transcriptomes, on average 5,902 genes were detected (counts larger than 1) in each cell, 41% (2,397) of which were lncRNAs, including 1,258 reference and 1,139 novel lncRNAs (Figure 3A). Furthermore, for both reference and novel lncRNA genes, the average number of expressed genes across T cells was significantly lower than that of protein-coding genes. More precisely, we found that an average of 5,596 protein-coding and 2,093 lncRNA genes were expressed in at least 25% of cells. In such a situation, novel lncRNAs exhibited a higher average expression number and expression rate (1,489 and 15.8%, 1,489/9,433) than did reference lncRNAs (604 and 1%, 604/59,489) (Figure 3B), suggesting that novel lncRNAs exhibited more enrichment than known lncRNAs in T cells in cancer. Moreover, we performed further analysis to investigate the specifically expressed lncRNAs in different tissues for each cancer type. In brief, for both CD4 and CD8 T cells of each cancer type, we identified 96 and 90 lncRNAs including 44 and 40 novel lncRNAs that expressed in tissue-specific pattern (Table S4). For example, some novel lncRNAs such as *XLOC-301694* and *XLOC-126527* were significantly expressed in CD4 T cells from tumor tissue of CRC (adjusted *P* value =3.17E-68 and 1.72E-64 respectively), while others such as *XLOC-302096* and *XLOC-502999* were significantly enriched in normal tissue and peripheral blood respectively (adjusted *P* value =9.18E-82 and 1.35E-44 respectively) (Table S4). Finally, we assessed the evolutionary conservation of these novel lncRNA transcripts and found that, on average, 61.2% have orthologous regions in the primate genomes, while only 3.4% mapped to mouse genome, suggesting the poor sequence conservation of these novel lncRNAs.

**Figure 3.**
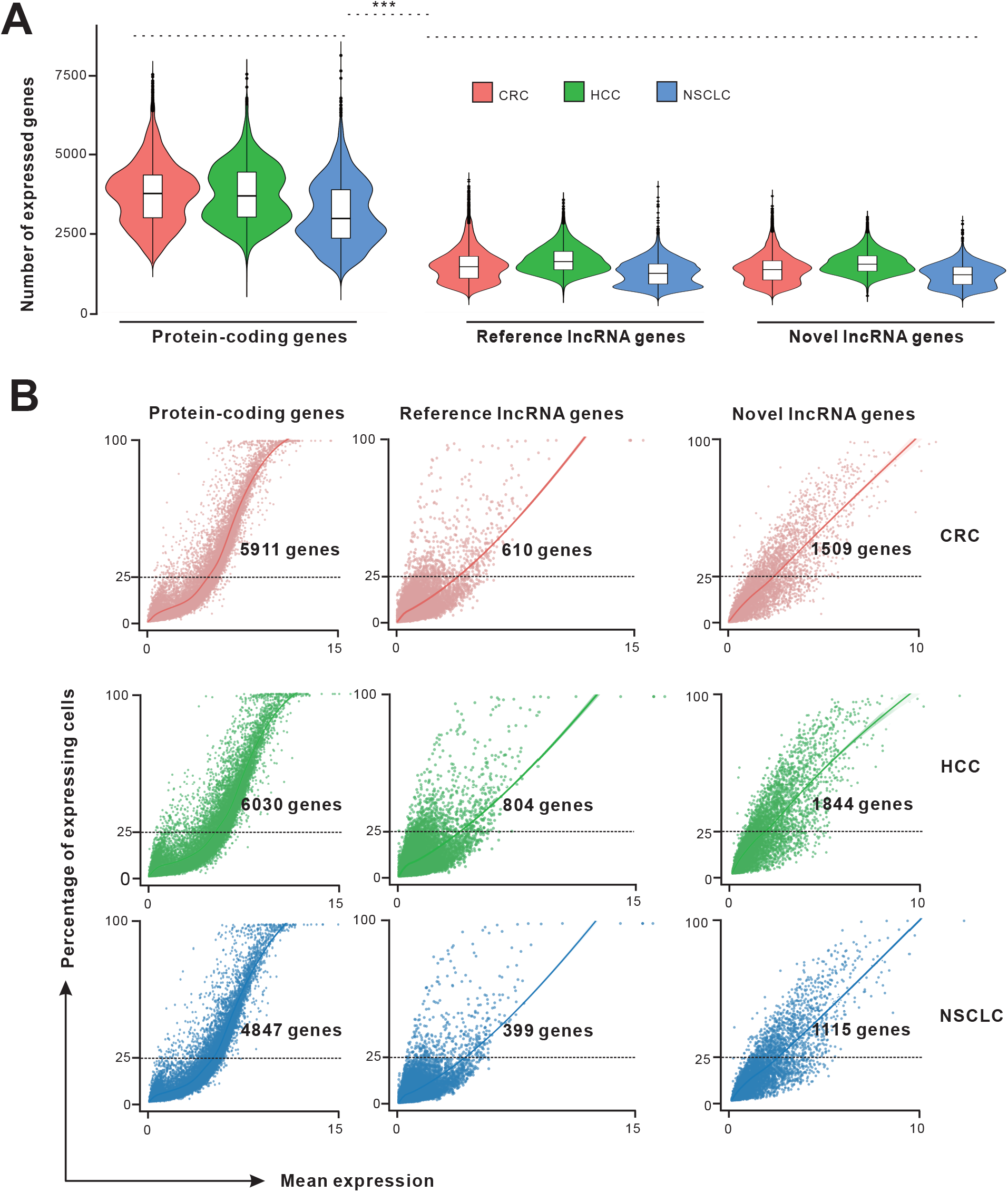
Characterization of lncRNA expression patterns at single-cell level. **A**. The number of protein-coding, reference lncRNA, and novel lncRNA genes expressed in T cells across three cancer types. *** indicates *P*-value < 0.001 (Wilcoxon rank sum test). **B**. The plots show the percentage of expressing cells against the mean expression level (logCounts) for protein-coding, reference lncRNA, and novel lncRNA genes across three cancer types. The numbers of genes that are expressed in at least 25% of cells are labelled.

### Identification of signature lncRNAs associated with T cell states in cancer immunity based on metacell maps

To explore signature lncRNAs associated with T cell states in cancer immunity, we used the MetaCell method [31] that partitioned the scRNA-seq datasets into disjointed and homogeneous cell groups (namely metacells) using the non-parametric *K*-nn graph algorithm. For the lowly and specifically expressed nature of lncRNA genes, metacells pooling together data from cells derived from the same transcriptional states could serve as building blocks for approximating the distributions of lncRNA gene expression and minimizing the technical variance and noise. After quality control, 19,572 cells with predefined cluster annotations and 21,205 genes including both protein-coding and lncRNA genes were retained and used for the following analyses. The expression tables of CD8 and CD4 T cells across three cancers were fed into the MetaCell pipeline separately, resulting in a detailed map of 43 and 65 metacells respectively (**Figure 4A-B** and Table S5-6).

**Figure 4.**
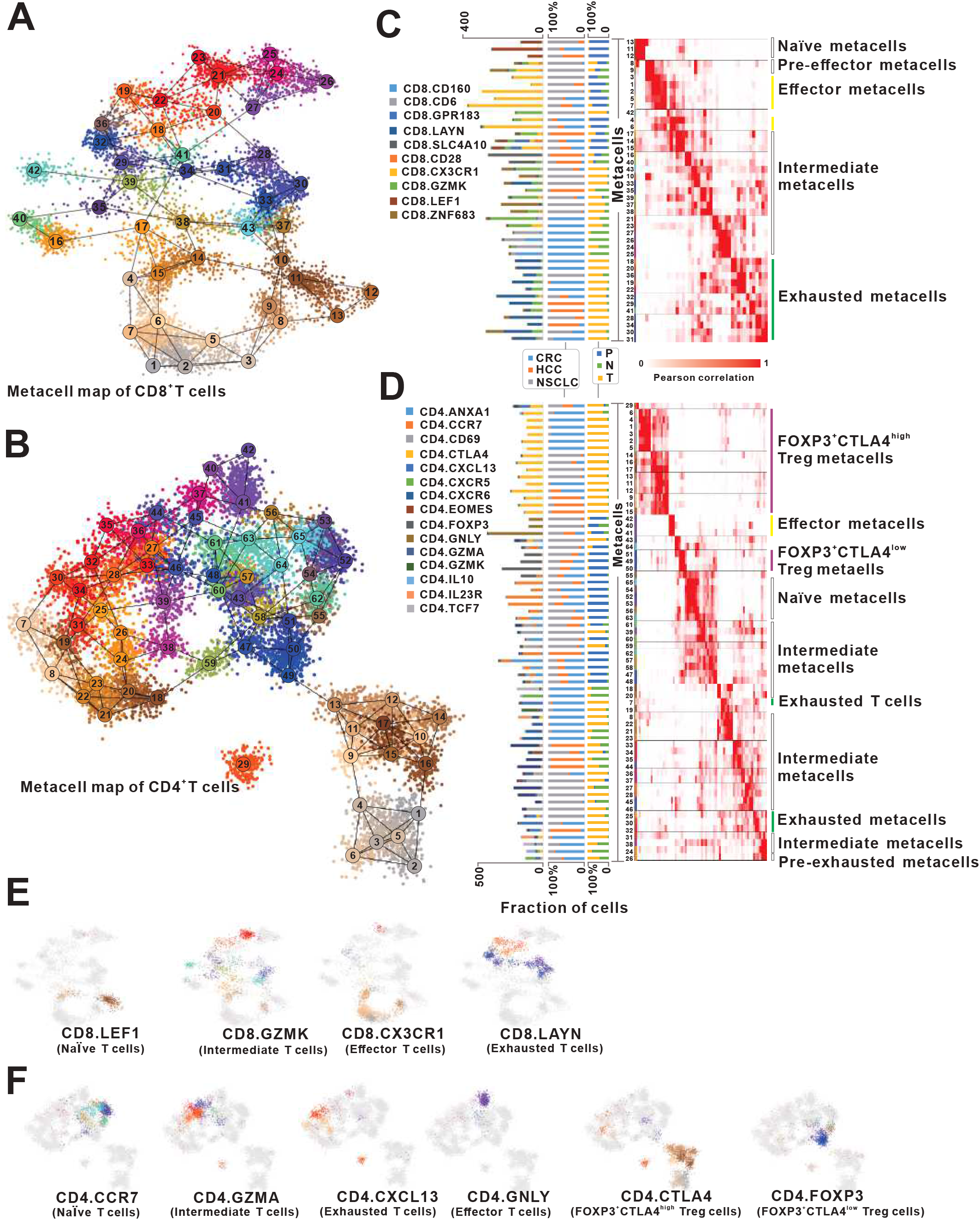
Characterization of T cell states based on 2D projection of T cells and the annotation of metacell maps. **A**. 2D projection of CD8 T cells from three cancer types into 43 metacells. **B**. 2D projection of CD4 T cells from three cancer types into 65 metacells. **C**, **D**. CD8 (**C**) and CD4 (**D**) metacells (rows) are ordered by groups and organized within each group. The first panel of the bar plot shows the number of cells of different clusters in each metacell. The second and third panel of the bar plots show the percentage of cells from different cancer types and tissues in each metacell respectively. Heatmaps show the confusion matrix (the pairwise similarities between metacells) for CD8 (**C**) and CD4 (**D**) metacells. The annotations of different metacell groups are shown on the right. **E**, **F**. 2D projections of the composition of CD8 (**E**) and CD4 (**F**) T cells from different clusters. P, peripheral blood; N, adjacent normal tissue; T, tumor tissue.

Based on the 2D projections (Figure 4A-B), predefined cell cluster annotations (Table S1), and the metacell similarity matrices (similarity among 43 or 65 metacells for CD8 or CD4 T cells respectively) (Figure S2A-B and Figure 4C-D), we organized the complex transcriptional landscape of CD8 into Naïve, effector/pre-effector, intermediated, and exhausted metacell groups and CD4 into Naïve, effector, intermediated, exhausted, and regulatory (including *FOXP3*^+^*CTLA4*^low^ and *FOXP3*^+^*CTLA4*^high^) metacell groups respectively (Figure 4C-D). To evaluate the composition of metacells, we mapped tissue- and cancer-specific patterns in all metacells and achieved results in accordance with previous studies [32–34] (Figure 4C-D and Figure S3-4). For example, exhausted metacells were preferentially enriched in tumors, while effector metacells were prevalent in peripheral blood. Although some metacells were enriched in different cancer types, they were organized into the same functional groups (Figure 4C-D). Notably, effector metacell groups (cytotoxic state) and exhausted metacell groups (dysfunctional state) were located in different directions in the metacell maps, while the diffuse border was observed between the intermediate and the cytotoxic or dysfunctional state (Figure 4E-F). These intermediate cells exhibited remarkable transcriptional heterogeneity indicating functional divergence of these cells (Figure 4E-F and Figure S3-4). The observed cluster distribution in both CD8 and CD4 metacell maps might suggest a relative transition from activation to exhaustion that began with Naïve cells, followed by intermediate cells (such as central memory (CM), effector memory (EM) and tissue resident memory (RM) cells) and ended with exhausted cells. Moreover, the CD4 metacell map revealed that Tregs were subdivided into *FOXP3*^+^*CTLA4*^low^ Tregs and *FOXP3*^+^*CTLA4*^high^ Tregs that were preferentially enriched in blood and tumors respectively (Figure 4D and 4F). These observations demonstrated that the diversity and dynamics of T cell states in cancer immune infiltrates could be controlled by complex and intricate gene regulatory mechanisms. Yet, the association between these cell states and lncRNAs was still poorly characterized, prompting us to subsequently investigate potential roles of lncRNA genes in T cells. Currently, the cell groups such as *FOXP3*^+^*CTLA4*^high^ Tregs and exhausted T cells expressing inhibitory receptors (e.g., *PDCD1* and *TIGIT*) have been used as therapeutic targets for anti-cancer immunotherapies, thus we focused on these cells in the following analyses.

To explore signature lncRNAs associated with effector T cells, exhausted T cells, and Tregs, we performed systematic analysis of these metacell groups based on well-defined anchor genes [39], such as the genes associated with CD8 effector functions (*CX3CR1, FGFBP2, GZMH* and *PRF1*) or with the CD8 exhausted state (*HAVCR2, LAG3, PDCD1, TIGIT* and *CTLA4*). As a result, 154 lncRNAs that were significantly correlated to the anchor genes were identified and were involved in a set of co-expressed gene modules, including effector, exhausted and Treg gene modules (**Figure 5A-B** and Table S7). Interestingly, a putative *CTLA4*^high^ Treg gene subset was observed in the Treg module, suggesting its specific functional roles in tumor-infiltrating Treg cells (Figure 5B). Overall, by combination analysis with the expression profile across metacell groups, we found 47 and 79 lncRNAs correlated with effector and exhausted states in CD8 and CD4 cells respectively and were designated as effector or exhausted signature lncRNAs (Figure 5C and Figure S5). Similarly, 49 lncRNAs were highly associated with Treg cells and were designated as Treg signature lncRNAs (Figure S5). Among these signature lncRNAs, 14 and 7 lncRNA genes were shared between CD8 and CD4 effector states and between CD8 and CD4 exhausted states respectively. 21 lncRNA genes associated with Tregs overlapped with those characteristics in the exhausted CD4 T cells (Table S7), indicating the presence of shared regulatory roles of these lncRNAs. In contrast, no signature lncRNA was shared between exhausted and effector states.

**Figure 5.**
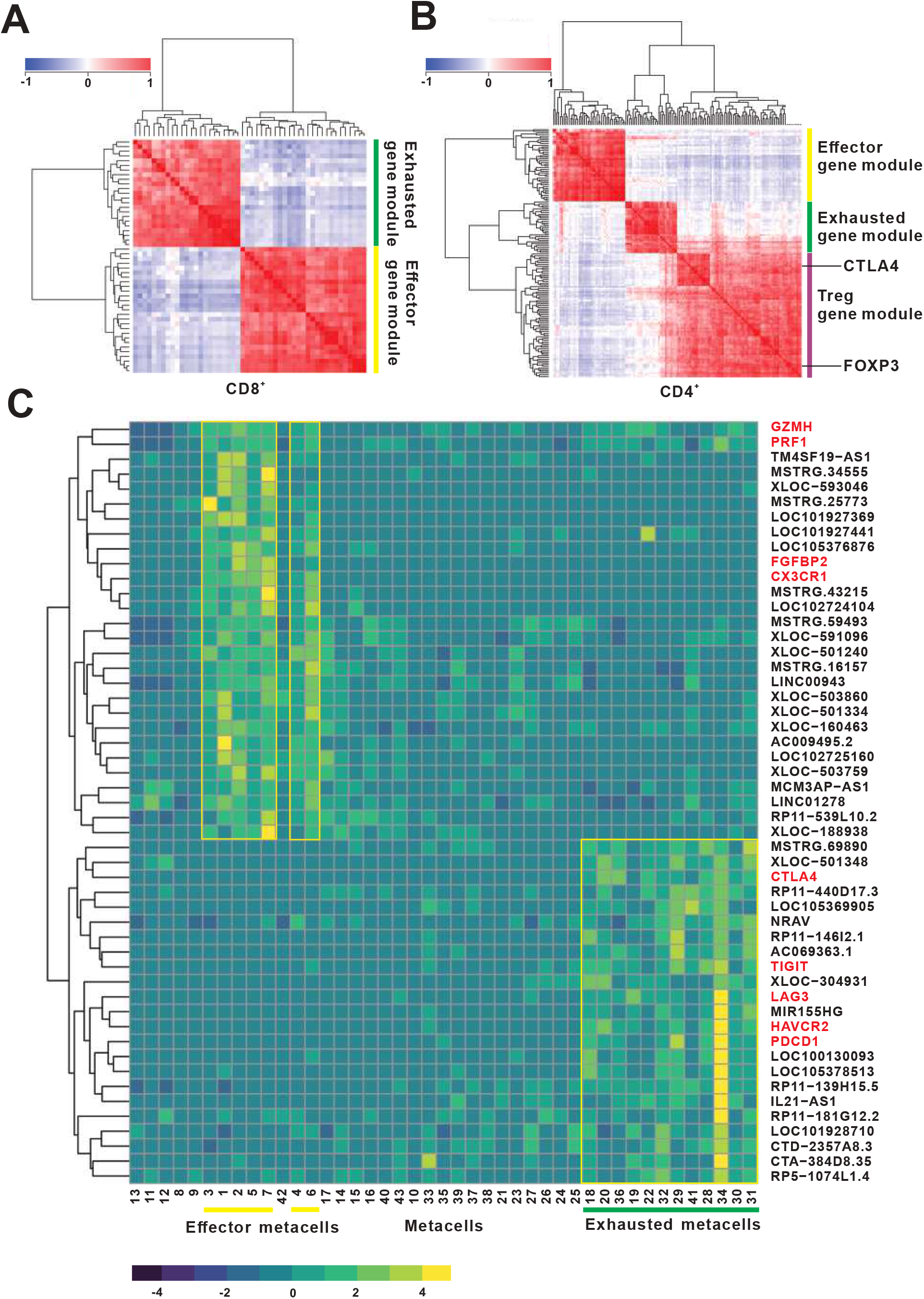
The correlation and expression analyses of signature lncRNAs associated with different T cell states. **A**, **B**. Gene-gene correlation heatmap for signature lncRNA and anchor genes in CD8 (A) and CD4 (B) T cells. The signature gene modules and two anchor genes (*CTLA4* and *FOXP3*) are labelled on the right. **C**. Expression of signature lncRNA and anchor genes across CD8 metacells. Metacells and metacell groups associated with effector and exhausted functions are shown on the bottom. The anchor genes are marked with red color on the right.

### Functional prediction of signature lncRNAs associated with T cell states based on co-expression network

To gain further insights into the functional roles of lncRNA in different T cell states in cancer, we built a coding-noncoding network (CNC), as we previously reported [40, 41], using linear correlation over all metacells. Applying this strategy, the functions of 54% (84/154) signature lncRNAs were annotated (Table S8). As expected, both CD8 and CD4 exhausted T cells have the functional enrichments of signature lncRNAs that were markedly different from effector CD8 or CD4 T cells, including regulation manners in immune system processes and several signalling pathways (**Figure 6A-B**). For example, exhausted signature lncRNAs were significantly enriched in immunoinhibitory functions such as negative regulation of immune response (adjusted *P*-value = 2.96e-14), negative regulation of T cell activation (adjusted *P*-value = 1.24e-06), and positive regulation of interleukin-10 biosynthetic process (adjusted *P*-value = 1.02e-18). In comparison, effector signature lncRNAs were enriched in cytotoxic programs such as T cell proliferation involved in immune response (adjusted *P*-value = 8.16e-09), positive regulation of cytokine secretion (adjusted *P*-value = 4.65e-05), and positive regulation of cytolysis (adjusted *P*-value = 1.59e-19) (Figure 6A-B and Table S9-10). These results consisted with the phenotypes of exhausted or effector states of T cells as described in previous studies [1, 32–34, 42]. In addition, the enriched functions of Treg signature lncRNAs were similar with those of CD4 exhausted signature lncRNAs involving multiple immunosuppressive programs (**Figure 6C** and Table S11), suggesting the shared regulatory roles of these lncRNAs in CD4 Tregs and exhausted CD4 T cells. Further analysis of the functions of co-signature lncRNAs that were shared between CD8 and CD4 exhausted or effector states, as well as between CD4 exhausted and Treg states (Figure S6), suggests that the signature lncRNAs might broadly participate in regulation of T cell functions within the human tumor microenvironment.

**Figure 6.**
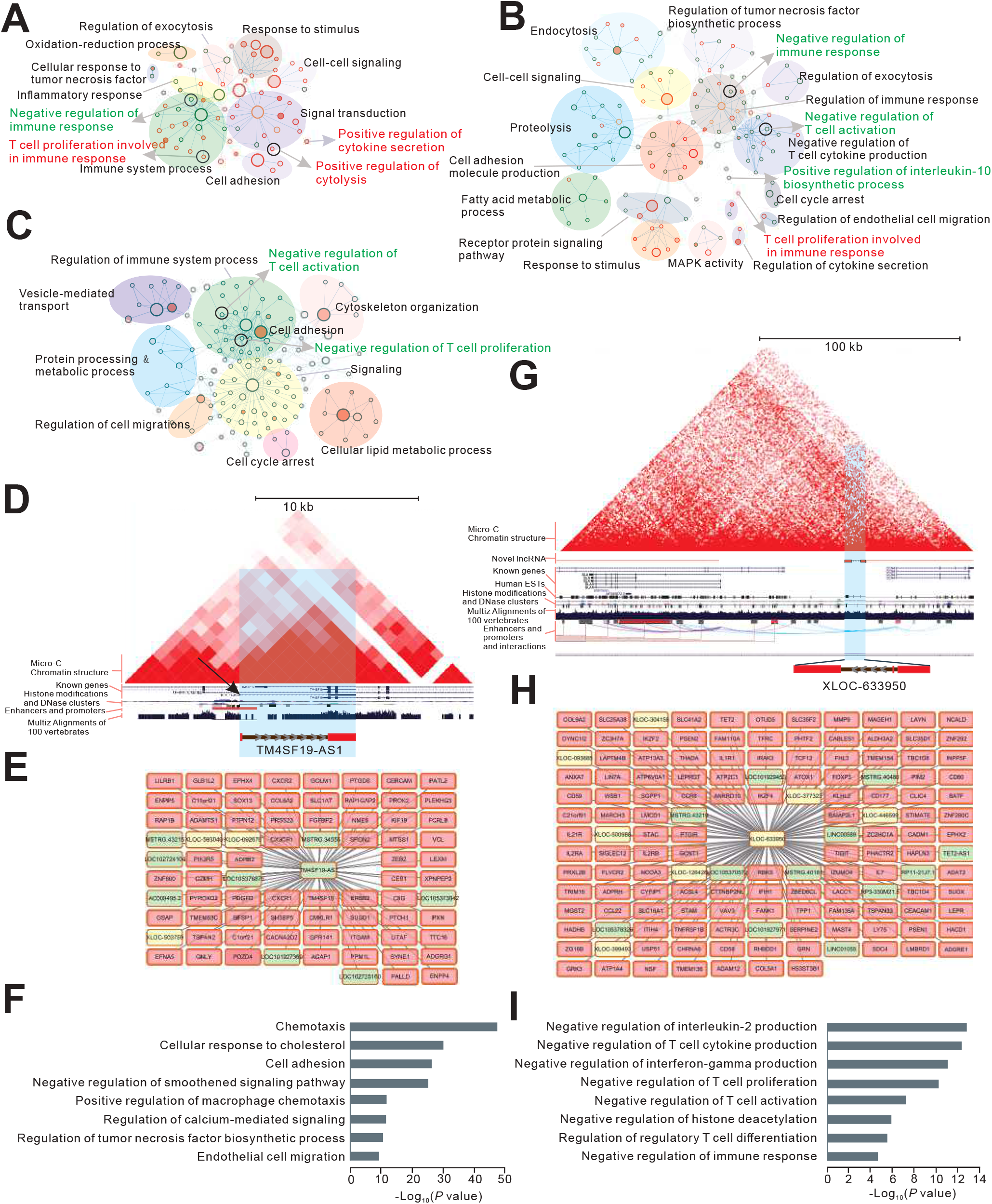
Functional annotation analyses of signature lncRNAs. **A**-**C**. Functional enrichment maps of CD8 effector/exhausted (**A**), CD4 effector/exhausted (**B**) and CD4 Treg (**C**) signature lncRNAs. The enriched gene sets from Gene Ontology based on the predicted functions of signature lncRNA genes are visualized by Cytoscape plugin Enrichment Map. Each node represents a gene set; size of the node is indicative of the number of genes and the color intensity reflects the level of significance. Effector signature gene sets are shown in red circles, exhausted or Treg ones in green and the common gene sets in orange. Maps are differently magnified for easier visualization. **D**-**F**. The genomic view (**D**), co-expressed genes (**E**) and functional annotations (**F**) of effector signature lncRNA *TM4SF19-AS1*. **G**-**I**. The genomic view (**G**), co-expressed genes (**H**) and functional annotations (**I**) of exhausted signature lncRNA *XLOC-633950* (novel). The genomic views are generated from UCSC genome browser. In (**E**) and (**H**), co-expressed protein-coding, reference lncRNA and novel lncRNA genes are colored by pink, light green and light yellow respectively.

For example, a known lncRNA *TM4SF19-AS1*, defined as a signature lncRNA for both CD8 effector and CD4 effector T cells and was transcribed in the antisense orientation to the *TM4SF19* gene, was co-expressed with 66 protein-coding and 11 lncRNA genes (Figure 6D-E). Of note, *TM4SF19-AS1* was highly correlated and located in the same topologically associated domain (TAD) with its host gene *TM4SF19* (Pearson correlation coefficient = 0.88) (Figure 6D), a member of the four-transmembrane L6 superfamily participating in various cellular processes including cell proliferation, motility, and cell adhesion [43–46]. Consistently, *TM4SF19-AS1* was significantly enriched in several effector T cell associated processes such as cellular response to cholesterol (adjusted *P*-value = 1.09e-30), cell adhesion (adjusted *P*-value = 5.25e-27) and regulation of tumor necrosis factor biosynthetic process (adjusted *P*-value = 3.75e-11) (Figure 6F). Interestingly, a recent study suggested that anti-tumor response of CD8 T cells could be enhanced by regulating cholesterol metabolism [47]. For another example, a novel lncRNA *XLOC-633950*, defined as a signature lncRNA for both CD4 exhausted T cells and Treg cells, was an intergenic gene and transcribed from the promoter-enhancer cluster region of the *SLA* and *CCN4* genes (Figure 6G). Furthermore, *XLOC-633950* as a novel gene, whose expression was supported by multiple expressed sequence tags (EST), was located in the same TAD with the *SLA* gene which acted as an inhibitor of antigen receptor signalling by negative regulation of positive selection and mitosis of T cells [48–51] (Figure 6G). In accordance with *SLA* functions, the functional enrichments of *XLOC-633950* according to its co-expressed protein-coding genes were mainly associated with immunoinhibitory processes, such as negative regulation of T cell cytokine production (adjusted *P*-value = 4.56e-13) and negative regulation of T cell proliferation and activation (adjusted *P*-value = 7.25e-11 and 5.85e-08 respectively) (Figure 6H-I). These results provided a starting point for future dissecting the mechanisms of signature lncRNAs.

## Discussion

Despite the obvious advantages, most scRNA-seq data was still limited in its ability to study lncRNAs, which were emerging as central players and key regulators in a number of biological processes such as anti-tumor immune response [52, 53]. In comparison with many scRNA-seq methods that amplified only the 3’ end of transcripts, the SMART-seq2 protocol could generate full-length cDNA from polyadenylated transcripts which results in data suitable for analysis of lncRNAs [30, 54]. In current study, we preformed systematic analyses of SMART-seq2 full-length scRNA-seq datasets and provided the first comprehensive atlas of lncRNA in T cells of human cancer.

Recently, Jiang *et al.* presented a comprehensive human lncRNA catalog (RefLnc) [38] containing 77,900 lncRNAs based on analysis of 14,166 polyA(+) RNA-Seq libraries and previous known annotations. Among the RefLnc lncRNAs, only 16% could be assembled and expressed in T cells. In addition, compared with bulk-seq data, scRNA-seq data could detected more known and novel transcripts. These observations suggested that despite the vast number of lncRNAs that have been identified using bulk-seq data [10, 12, 26, 38, 55], the catalog of human lncRNAs is still far from being complete at single-cell resolution, due to their low and cell-specific expression patterns. Based on the cell-pooling strategy and more than 20,000 scRNA-seq libraries from 31 patients across three cancer types, we identified 9,433 previously non-annotated lncRNAs. These results significantly expand the current lncRNA catalog and enable us to carry out in-deep analysis of the T cell context-specific lncRNA transcriptome. Notably, all the scRNA-seq data used in current study was generated by sequencing the polyadenylated (ployA) transcriptome, in which non polyadenylated lncRNAs were absent.

Several previous studies have applied full-length scRNA-seq to unleash tumor infiltrating lymphocytes in HCC [34], NSCLC [32], and CRC[33], providing a deep understanding of the immune landscape of T cells in cancer. Nevertheless, the physiological function of lncRNAs in different T cell states during the cancer immune response remains elusive. Although the abundance of lncRNA was relatively low and hard to distinguish from technical noise in single T cells, pooling the transcripts from multiple cells that are derived from the same cell state allows more accurate quantification of lncRNAs, making it feasible to explore their signatures and putative regulatory mechanisms associated with T cell states in cancer immunity. Based on such partitioning and pooling strategies, we used the MetaCell method to identify homogeneous T cell groups from scRNA-seq data and derived a detailed map of 43 and 65 metacells for CD8 and CD4 T cells respectively. These metacells with higher homogeneity, allowed a more accurate quantification of lncRNAs as well as identification of T cell differentiation gradients. For example, we observed 7 metacells involved in CD8 effector cell cluster, which might reflect the transcriptional heterogeneity in this cluster (Figure 4C). The roles of lncRNAs in these different subsets (metacells) of CD8 effector T cells need further investigation. While MetaCell was not designed to perform single-cell lncRNA analysis, the MetaCell partitioning algorithm facilitated robust cell grouping of scRNA-seq data which enabled us to study lncRNAs more accurately.

According to the metacell maps (Figure 4E-F), in contrast to the pool of intermediate T cells with diffuse borders with other cell states, a discrete pool of effector T cells, exhausted T cells and Tregs were observed that show clear gaps among them, thus facilitating unbiased analysis of signature lncRNAs in these cell states. In total, the 154 signature lncRNAs were obtained providing a useful reference lncRNA resource to further investigate their functions in T cell mediated cancer immunity. Since lncRNAs generally interact with protein-coding genes, and highly correlated genes generally have similar functions, the putative functions of these signature lncRNAs could be predicted by the co-expressed coding genes. Therefore, by constructing the ‘two color’ co-expression network in which both coding and lncRNA genes were involved, the functions of 84 signature lncRNAs were annotated. Some lncRNAs were genomically co-located with their host genes, that revealed the complicated regulation mechanisms of lncRNAs in cancer immunity. For example, as described above, *TM4SF19-AS1* was both co-expressed and co-located with their host gene *TM4SF19*, whose family has functions in various biological processes including cell proliferation and adhesion that are consistent with the characteristics of effector T cells [43–46].

In summary, the current study provides the first comprehensive catalog and the functional repertoires of lncRNAs in human cancer T cells. Although the expression pattern and exact mechanisms of these signature lncRNAs in regulating T cell states needs further experimental validation, we provide the groundwork for future studies to investigate the functional mechanisms of lncRNAs in the T cell mediated cancer immunity, especially in two of the essential states of T cells: effector state and exhausted state. These signature lncRNAs of CD8 exhausted T cells and tumor Tregs, may serve as new targets for novel cancer-immune biomarker development and cancer immunotherapies.

## Materials and methods

### Full-length scRNA-seq and bulk RNA-seq datasets from cancer patients

Raw sequencing data of three compendium datasets used in the current study were authorized by the European Genome-phenome Archive (EGA) and obtained from the EGA database under study accession id: EGAS00001002791, EGAS00001002430, and EGAS00001002072. The CRC scRNA-seq dataset (EGAS00001002791) contains the raw data of 11,138 single T cells isolated from different tissues (peripheral blood, adjacent normal and tumor tissues) of 12 CRC patients [33]. The NSCLC scRNA-seq dataset (EGAS00001002430) contains the raw data of 12,346 single T cells from 14 NSCLC patients [32]. The HCC scRNA-seq dataset (EGAS00001002072) contains the raw data of 5,063 single T cells from 6 HCC patients [34]. All the data were generated by Illumina HiSeq 2500 sequencer with 100 bp pair-end reads or Illumina Hiseq 4000 sequencer with 150 bp pair-end reads. The cells from HCC patient P1202 (TCRs could not be assembled in those cells) were not analyzed in the current study. After preliminary filtration, 24,075 T cells with at least one pair of TCR *alpha-beta* chain were retained. The bulk RNA-seq data of five tumor samples from HCC patients were obtained from HCC dataset.

According to the cell annotations from original papers [32–34], these T cells were classified into different subtypes (Figure S1A and Table S1). PTC, NTC, and TTC represent CD3^+^CD8^+^ T cells that were isolated from peripheral blood, adjacent normal, and tumor tissues respectively. The PTH, NTH, and TTH represent CD3^+^CD4^+^CD25^low^ T cells that were isolated from the three tissues. PTR, NTR, and TTR represent CD3^+^CD4^+^CD25^high^ T cells that were isolated from the three tissues.

### Reads mapping and transcripts assembly

Clean reads from each T cell were mapped to the human reference genome (version hg38/GRCh38) using STAR aligner (v2.7.1) [56] with the *twopassMode* set as Basic. The bam files of T cells from each cell-type of each patient were merged using SAMtools merge [57]. StringTie (v2.0.3) [35] was used to assemble transcripts based on genomic read alignments. Assembled transcripts of all cell-types across all patients were merged together using the Cuffmerge utility of Cufflinks package [58].

### Comparison with reference gene annotation

For reference gene annotation, lncRNA genes were collected from RefLnc [38] and other genes were collected from GENCODE v31 [59]. According to the “class code” information outputted by Cuffcompare, the merged assembly was classified into four categories by comparison with the reference gene annotation, including known coding genes, known lncRNA genes, potentially novel genes (class code is “i, x, u”), and others.

### Identification of novel lncRNAs

Based on the potentially novel gene catalog derived from single-cell data, we developed a custom pipeline for identification of reliable novel lncRNAs including the following steps: (1) transcripts that are no shorter than 200 nt and have more than one exon were selected for downstream analysis (for intergenic transcripts, at least 1 kb away from known protein-coding genes); (2) CPC (Coding Potential Calculator) [36] and CNCI (Coding Noncoding Index) [37] software were used to evaluate the protein-coding potential of transcripts, and transcripts that were reported to lack coding potential by both CPC and CNCI were regarded as candidate noncoding transcripts; (3) The remaining transcripts that were assembled and have the same intron chain of at least two cell-types were retained as the final novel lncRNA catalog. The final lncRNA catalog was obtained by combining the reference lncRNA and novel lncRNA genes directly. The UCSC liftOver tool (http://genome.ucsc.edu/cgi-bin/hgLiftOver?hgsid=806106955_h2xhcK2iPRI7SiMkxkB41I2mwF9O) was used to identify the orthologous locations of human novel lncRNAs in the mouse genome and in primates such as chimpanzee and gorilla, with the parameters: Minimum ratio of bases that must remap = 0.1 and Min ratio of alignment blocks or exons that must map =0.5.

### Experimental validation of novel lncRNAs

Three CRC patients were enrolled at Shenzhen People’s Hospital. The informed consent forms were provided by patients. The current study was approved by Medical Ethics Committee of Shenzhen People’s Hospital. The clinical characteristics of three patients are summarized in Table S2. Peripheral blood samples from three patients were obtained and treated with anticoagulation. Peripheral blood mononuclear cells (PBMCs) were extracted by Ficoll-Paque Plus (GE Healthcare, Sweden, 17144003). Then, CD8^+^ and CD4^+^ T cells were separated by immunomagnetic beads (Meltenyi Biotec, Germany, 130045101, 130045101). The separation efficiency was verified by flow cytometry. The sorted cells were dissolved in Trizol Reagent (Ambion, USA, 15596026) for RNA extraction according to the manufacture’s protocol. cDNA was synthesized by PrimerScript RT reagent kit (Takara, Japan, AHG1552A). We chose 50 novel lncRNAs to perform experimental validation according to the following criteria: (1) highly expressed in either CD8 or CD4 T cells; (2) reconstructed in at least ten subsets with complete match; (3) uniquely mapped to human genome. For each lncRNA, at least two pairs of primers for qRT-PCR were designed using NCBI Primer-BLAST (https://www.ncbi.nlm.nih.gov/tools/primer-blast). In order to ensure the specificity of primers, UCSC InSilicon PCR (http://genome.ucsc.edu/cgi-bin/hgPcr) was used to compare the primer pairs with human genome (hg38). Some primer pairs were specifically designed to span splicing sites (exon junctions). QRT-PCR were performed with SYBR Green master mix (Takara, Japan) on an ABI StepOnePlus (Applied Biosystems, USA). *GAPDH* as housekeeping gene was used as positive control. For each lncRNA, we selected one primer pair product of qRT-PCR for Sanger sequencing.

### Quality control (QC) and normalization

We calculated the read counts and transcripts per million (TPM) values using pseudoalignment of scRNA-seq reads to both protein-coding and lncRNA transcriptomes, as implemented in Kallisto (v0.46.0) [60] with default parameters, and summarized expression levels from the transcript level to the gene level.

Low-quality and doublet cells were removed if the number of expressed genes (counts of more than 1) was fewer than 2000 or higher than the medians of all cells plus 3 × the median absolute deviation, respectively. Moreover, the cells with the proportion of reads mapped to mitochondrial genes was larger than 10% were discarded. Genes with average counts of more than 1 and expressed in at least 1% of cells for each type of cancer were retained. The combined count tables from all T cells passing the above filtration were normalized using a pooling and deconvolution method implemented in the R package named *computeSumFactors* [61] with the sizes ranged from 80, 100, 120 to 140. According to the assumption that most genes were not differentially expressed, normalization was performed within each predefined cluster separately to compute cell size factors. The cell size factors were rescaled by normalization among clusters. Finally, the counts for each cell were normalized by dividing the cell counts by the cell size factor.

### MetaCell modeling

The MetaCell method [31], that partitioned the scRNA-seq dataset into disjointed and homogeneous cell groups (metacells) using the *K*-nn graph algorithm, was performed for both the CD8 and CD4 T cells independently. We first removed specific mitochondrial genes (annotated with the prefix “MT-”), that typically mark cells as being stressed or dying, rather than cellular identity. Based on the count matrices of both protein-coding and lncRNA genes, feature genes whose scaled variance (variance/mean on down-sampled matrices) exceeded 0.08 were selected and used to compute cell-to-cell similarity using Pearson correlations. According to the cell-to-cell similarity matrices, two balanced *K*-nn similarity graphs for CD8 and CD4 T cells were constructed using the parameter *K*=100 (the number of neighbors for each cell was limited by *K*). Next, we performed the resampling procedures (resampling 75% of the cells in each iteration with 500 iterations) and co-clustering graph construction (the minimal cluster size was 50). Finally, the graphs of metacells (and the cells belonging to them) were projected into 2D spaces to explore the similarities between cells and metacells.

### Annotation of metacells

Annotation of metacells was performed based on the metacell confusion matrix and predefined cluster annotations (File S1) of T cells involved in the metacells. Briefly, we first created a hierarchical clustering of metacells according to the number of similarity relationships between their cells. Next, we generated clusters of metacells as confusion matrices based on the hierarchy results, then annotated these clusters according to the annotations of T cells.

### Defining signature lncRNAs associated with T cell states

To identify signature lncRNAs associated with effector and exhausted T cells as well as Tregs, as described in recent study [39], we adopted the anchor approach by identifying the lncRNAs that were significantly correlated to well-defined anchor genes, based on metacells’ log enrichment scores (*lfp* values calculated by MetaCell method). The lncRNAs that significantly correlated with anchor genes (adjusted *P*-value <0.01 and ranked in the top 0.05 percentile for each anchor gene) were regarded as signature lncRNAs. The anchor genes were defined as follows: the anchor genes of CD8 exhausted T cells included *HAVCR2, LAG3, PDCD1, TIGIT*, and *CTLA4*; the anchor genes of CD8 effector T cells included *CX3CR1, FGFBP2, GZMH* and *PRF1*; genes associated with Tregs included *FOXP3*; the anchor genes of CD4 exhausted T cells included *CXCL13, PDCD1, HAVCR2, TIGIT*, and *CTLA4*; genes associated with CD4 effector T cells included *GNLY, GZMB, GZMH, PRF1*, and *NKG7*.

### Function prediction of signature lncRNAs based on co-expression network

Based on *lfp* values of both lncRNA and protein-coding genes across all metacells, we used a custom pipeline for large-scale prediction of signature lncRNA functions by constructing the coding-lncRNA gene co-expression network [40, 41]. Briefly, genes with log enrichment scores ranked in the top 75% of each metacell were retained. Then, *P*-values of Pearson correlation coefficients for each gene pair were calculated based on the Fisher’s asymptotic test using the *WGCNA* package of R. *P*-values were adjusted based on the Bonferroni multiple test correction using the *multtest* package of R. The gene pairs with an adjusted *P*-value < 0.01, Pearson correlation coefficient > 0.7, and ranked in the top 5% for each gene were involved in co-expression network.

Based on the co-expression network, lncRNA functions were predicted using module- and hub-based methods. Specifically, the Markov cluster algorithm was adopted to identify co-expressed modules [40]. For each module, if the known genes were significantly enriched for at least one Gene Ontology (GO) term, the functions of the lncRNAs involved in the module were assigned as the same ones. For hub-based method, the functions of a hub lncRNA (node degree > 10) were assigned, if its immediate neighboring genes were significantly enriched for at least one GO term.

## Data availability

All the novel lncRNA genes identified in current study and their expression files are available in the NONCODE database (http://www.noncode.org/download.php).

## Authors’ contributions

HL, DB, JD, YZ and FL conceptualized and designed the study. HL and DB led the data analysis. HL performed the study and interpreted data. LJS performed experimental validation. YL collected the clinical samples and prepared the experimental materials. LS optimized the CNCI algorithm. WY, CW, XY and JW collected the data and performed T cell annotations. HL wrote the manuscript. HL, YZ and FL revised the manuscript. JD, YZ and FL supervised the project. All authors read and approved the final manuscript.

## Acknowledgments

This work was supported by the Science and Technology Project of Shenzhen (No. JHZ20170310090257380, JCYJ20170413092711058, JCYJ20170307095822325), China Postdoctoral Science Foundation (No. 2019M663369), Natural Science Foundation of Shenzhen (20190727160324164), and National Natural Science Foundation of China (31970636). We thank Dr. Lei Zhang and Yao He at Peking University for assistance with raw data download. The data analysis was performed on AWS (China region) and we thank Mr Hansen Huang, Chao Wu and Fei Shi for providing the technical service and support.

## Supplementary material

**Figure S1 The statistics of T cell data analysis**

**A**. The number of cells in different subsets across all patients from three cancer types. **B**, **C**. The number **(B)** and the ratio **(C)** of uniquely mapped read pairs of T cell sequencing data. **D**. The number of splices of mapping results. **E**. The different strategies used to explore the best way to obtain novel transcripts. **F**. The number of assembled transcripts in each subset. PTC, CD8^+^ cytotoxic T cells from peripheral blood; TTC, CD8^+^ cytotoxic T cells from tumor tissue; NTC, CD8^+^ cytotoxic T cells from adjacent normal tissue; PTH, CD4^+^CD25^−^ cells from peripheral blood; TTH, CD4^+^CD25^−^ cells from tumor tissue; NTH, CD4^+^CD25^−^ cells from adjacent normal tissue; PTR, CD4^+^CD25^hi^ cells from peripheral blood; TTR, CD4^+^CD25^hi^ cells from tumor tissue; NTR, CD4^+^CD25^hi^ cells from adjacent normal tissue; PTY, CD4^+^CD25^int^ cells from peripheral blood; TTY, CD4^+^CD25^int^ cells from tumor tissue; NTY, CD4^+^CD25^int^ cells from adjacent normal tissue; PPQ, CD4^+^ T cells from peripheral blood; TPQ, CD4^+^ T cells from tumor tissue; NPQ, CD4^+^ T cells from adjacent normal tissue; CRC, colorectal cancer; HCC, hepatocellular carcinoma; NSCLC, non-small-cell lung cancer.

**Figure S2 The cluster hierarchy of metacells**

**A**, **B**. The cluster hierarchy of CD8 (**A**) and CD4 (**B**) metacells. Subtrees in blue, sibling subtrees in gray. The metacells are colored and labelled on bottom.

**Figure S3 2D projections of CD8 T cells**

**A**, **B**. The composition of CD8 T cells from different clusters (**A**) and cancer types (**B**). Metacells and the cells involved in them are marked by different colors. The number of cells within each cluster is shown in brackets.

**Figure S4 2D projections of CD4 T cells**

**A**, **B**. The composition of CD4 T cells from different clusters (**A**) and cancer types (**B**). Metacells and the cells involved in them are marked by different colors. The number of cells within each cluster is shown in brackets.

**Figure S5 Expression of signature lncRNA and anchor genes across CD4 metacells**

Metacells and metacell groups associated with effector, exhausted and Treg functions are shown on the bottom. The anchor genes are marked with red color on the right.

**Figure S6 Functional enrichment maps of shared signature lncRNAs**

**A**-**C**. Functional enrichment maps of shared signature lncRNAs between CD8 effector and CD4 effector function (**A**), between CD8 exhausted and CD4 exhausted function (**B**) and between CD4 exhausted and CD4 Treg function (**C**). Each node represents a gene set; size of the node is indicative of the number of genes and the color intensity reflects the level of significance. Maps are differently magnified for easier visualization.

**Table S1 The basic information of single T cell data**

**Table S2 Clinical characteristics of three cancer patients**

**Table S3 The list of novel lncRNAs successfully validated by Sanger sequencing**

**Table S4 The list of specific-expressed lncRNAs**

**Table S5 The composition of CD8 metacells**

**Table S6 The composition of CD4 metacells**

**Table S7 The list of signature lncRNAs**

**Table S8 Functional annotations of 84 signature lncRNAs**

**Table S9 Functional enrichment results of CD8 effector/exhausted signature lncRNAs**

**Table S10 Functional enrichment results of CD4 effector/exhausted signature lncRNAs**

**Table S11 Functional enrichment results of CD4 Treg signature lncRNAs**

Supplementary Table1-8 are Excel format, and Supplementary Table9-11 are Word format.

