## Supplementary material for "Single-cell Long Non-coding RNA Landscape of T Cells in Human Cancer Immunity": Table S9.docx

**Table S9 Functional enrichment results of CD8 effector/exhausted signature lncRNAs**

| **GO ID** | **GO name** | **Adjusted *P*-value** | **Status** |
| --- | --- | --- | --- |
| GO:0071397 | cellular response to cholesterol | 1.35E-26 | Effector T cells |
| GO:0007224 | smoothened signaling pathway | 9.89E-26 | Effector T cells |
| GO:0006546 | glycine catabolic process | 1.22E-25 | Exhausted T cells |
| GO:0045879 | negative regulation of smoothened signaling pathway | 1.05E-21 | Effector T cells |
| GO:0045919 | positive regulation of cytolysis | 1.59E-19 | Effector T cells |
| GO:0021801 | cerebral cortex radial glia guided migration | 8.66E-17 | Effector T cells |
| GO:0006887 | exocytosis | 2.19E-16 | Exhausted T cells |
| GO:0032402 | melanosome transport | 3.80E-16 | Exhausted T cells |
| GO:0010573 | vascular endothelial growth factor production | 4.75E-16 | Effector T cells |
| GO:0040015 | negative regulation of multicellular organism growth | 4.59E-15 | Effector T cells |
| GO:0050777 | negative regulation of immune response | 2.96E-14 | Exhausted T cells |
| GO:0043123 | positive regulation of I-kappaB kinase/NF-kappaB cascade | 1.26E-13 | Common |
| GO:0045921 | positive regulation of exocytosis | 4.15E-13 | Exhausted T cells |
| GO:0070528 | protein kinase C signaling cascade | 1.16E-12 | Effector T cells |
| GO:0002548 | monocyte chemotaxis | 2.46E-12 | Exhausted T cells |
| GO:0032945 | negative regulation of mononuclear cell proliferation | 1.05E-11 | Effector T cells |
| GO:0032400 | melanosome localization | 7.20E-11 | Exhausted T cells |
| GO:0010759 | positive regulation of macrophage chemotaxis | 9.23E-11 | Effector T cells |
| GO:0002250 | adaptive immune response | 1.43E-10 | Exhausted T cells |
| GO:0042534 | regulation of tumor necrosis factor biosynthetic process | 2.75E-10 | Effector T cells |
| GO:0055114 | oxidation-reduction process | 3.33E-10 | Exhausted T cells |
| GO:0035025 | positive regulation of Rho protein signal transduction | 5.52E-10 | Effector T cells |
| GO:0007186 | G-protein coupled receptor protein signaling pathway | 9.33E-10 | Effector T cells |
| GO:0048245 | eosinophil chemotaxis | 9.62E-10 | Exhausted T cells |
| GO:0006954 | inflammatory response | 1.18E-09 | Exhausted T cells |
| GO:0014063 | negative regulation of serotonin secretion | 1.31E-09 | Effector T cells |
| GO:0007229 | integrin-mediated signaling pathway | 1.35E-09 | Effector T cells |
| GO:0045077 | negative regulation of interferon-gamma biosynthetic process | 2.61E-09 | Effector T cells |
| GO:0045636 | positive regulation of melanocyte differentiation | 2.74E-09 | Effector T cells |
| GO:0034699 | response to luteinizing hormone stimulus | 5.74E-09 | Effector T cells |
| GO:0031295 | T cell costimulation | 5.76E-09 | Exhausted T cells |
| GO:0002309 | T cell proliferation involved in immune response | 8.16E-09 | Effector T cells |
| GO:0010875 | positive regulation of cholesterol efflux | 1.00E-08 | Effector T cells |
| GO:0035108 | limb morphogenesis | 1.00E-08 | Effector T cells |
| GO:0008589 | regulation of smoothened signaling pathway | 1.98E-08 | Effector T cells |
| GO:0030326 | embryonic limb morphogenesis | 2.04E-08 | Effector T cells |
| GO:0043542 | endothelial cell migration | 2.29E-08 | Effector T cells |
| GO:0060318 | definitive erythrocyte differentiation | 2.86E-08 | Effector T cells |
| GO:0048247 | lymphocyte chemotaxis | 9.50E-08 | Exhausted T cells |
| GO:0007155 | cell adhesion | 1.16E-07 | Effector T cells |
| GO:0002740 | negative regulation of cytokine secretion involved in immune response | 1.18E-07 | Effector T cells |
| GO:0042536 | negative regulation of tumor necrosis factor biosynthetic process | 2.05E-07 | Effector T cells |
| GO:0032609 | interferon-gamma production | 2.05E-07 | Effector T cells |
| GO:0045428 | regulation of nitric oxide biosynthetic process | 2.57E-07 | Effector T cells |
| GO:0045785 | positive regulation of cell adhesion | 2.58E-07 | Effector T cells |
| GO:0043306 | positive regulation of mast cell degranulation | 3.39E-07 | Effector T cells |
| GO:0010628 | positive regulation of gene expression | 3.46E-07 | Exhausted T cells |
| GO:0032354 | response to follicle-stimulating hormone stimulus | 3.98E-07 | Effector T cells |
| GO:0047484 | regulation of response to osmotic stress | 4.87E-07 | Effector T cells |
| GO:0007165 | signal transduction | 5.27E-07 | Common |
| GO:0002376 | immune system process | 6.42E-07 | Exhausted T cells |
| GO:0006437 | tyrosyl-tRNA aminoacylation | 6.53E-07 | Exhausted T cells |
| GO:0051092 | positive regulation of NF-kappaB transcription factor activity | 7.63E-07 | Exhausted T cells |
| GO:0034695 | response to prostaglandin E stimulus | 1.19E-06 | Effector T cells |
| GO:0045668 | negative regulation of osteoblast differentiation | 1.71E-06 | Effector T cells |
| GO:0032695 | negative regulation of interleukin-12 production | 2.87E-06 | Effector T cells |
| GO:0043615 | astrocyte cell migration | 3.40E-06 | Exhausted T cells |
| GO:0032691 | negative regulation of interleukin-1 beta production | 4.59E-06 | Effector T cells |
| GO:0050764 | regulation of phagocytosis | 4.89E-06 | Effector T cells |
| GO:0048489 | synaptic vesicle transport | 4.92E-06 | Exhausted T cells |
| GO:0015914 | phospholipid transport | 5.20E-06 | Effector T cells |
| GO:0001915 | negative regulation of T cell mediated cytotoxicity | 6.36E-06 | Exhausted T cells |
| GO:0060907 | positive regulation of macrophage cytokine production | 6.48E-06 | Effector T cells |
| GO:0030890 | positive regulation of B cell proliferation | 6.98E-06 | Exhausted T cells |
| GO:0051926 | negative regulation of calcium ion transport | 7.23E-06 | Effector T cells |
| GO:0016485 | protein processing | 7.47E-06 | Effector T cells |
| GO:0018126 | protein hydroxylation | 8.19E-06 | Exhausted T cells |
| GO:0045806 | negative regulation of endocytosis | 8.19E-06 | Effector T cells |
| GO:0071347 | cellular response to interleukin-1 | 9.46E-06 | Exhausted T cells |
| GO:0031623 | receptor internalization | 1.02E-05 | Effector T cells |
| GO:0043308 | eosinophil degranulation | 1.60E-05 | Exhausted T cells |
| GO:0002768 | immune response-regulating cell surface receptor signaling pathway | 1.71E-05 | Effector T cells |
| GO:0071346 | cellular response to interferon-gamma | 1.92E-05 | Exhausted T cells |
| GO:0007167 | enzyme linked receptor protein signaling pathway | 2.43E-05 | Effector T cells |
| GO:0007267 | cell-cell signaling | 2.54E-05 | Effector T cells |
| GO:0002767 | immune response-inhibiting cell surface receptor signaling pathway | 2.69E-05 | Effector T cells |
| GO:0051271 | negative regulation of cellular component movement | 3.29E-05 | Effector T cells |
| GO:0009615 | response to virus | 3.29E-05 | Effector T cells |
| GO:0006911 | phagocytosis, engulfment | 3.63E-05 | Effector T cells |
| GO:0060347 | heart trabecula formation | 3.92E-05 | Effector T cells |
| GO:0071621 | granulocyte chemotaxis | 4.38E-05 | Exhausted T cells |
| GO:0050715 | positive regulation of cytokine secretion | 4.65E-05 | Effector T cells |
| GO:0014808 | release of sequestered calcium ion into cytosol by sarcoplasmic reticulum | 6.54E-05 | Exhausted T cells |
| GO:0050901 | leukocyte tethering or rolling | 6.79E-05 | Effector T cells |
| GO:0070098 | chemokine-mediated signaling pathway | 7.42E-05 | Effector T cells |
| GO:0002430 | complement receptor mediated signaling pathway | 8.29E-05 | Effector T cells |
| GO:0007200 | activation of phospholipase C activity by G-protein coupled receptor protein signaling pathway coupled to IP3 second messenger | 8.32E-05 | Effector T cells |
| GO:0045088 | regulation of innate immune response | 8.81E-05 | Effector T cells |
| GO:0071356 | cellular response to tumor necrosis factor | 8.88E-05 | Exhausted T cells |
| GO:0007266 | Rho protein signal transduction | 9.06E-05 | Effector T cells |
