## Supplementary material for "Single-cell Long Non-coding RNA Landscape of T Cells in Human Cancer Immunity": Table S10.docx

**Table S10 Functional enrichment results of CD4 effector/exhausted signature lncRNAs**

| **GO ID** | **GO name** | **Adjusted *P*-value** | **Status** |
| --- | --- | --- | --- |
| GO:0071397 | cellular response to cholesterol | 1.21E-26 | Effector T cells |
| GO:0007220 | Notch receptor processing | 3.58E-26 | Exhausted T cells |
| GO:0007224 | smoothened signaling pathway | 8.90E-26 | Effector T cells |
| GO:0006546 | glycine catabolic process | 9.18E-26 | Exhausted T cells |
| GO:0032468 | Golgi calcium ion homeostasis | 4.67E-25 | Exhausted T cells |
| GO:0032472 | Golgi calcium ion transport | 5.58E-24 | Exhausted T cells |
| GO:0015718 | monocarboxylic acid transport | 5.94E-23 | Exhausted T cells |
| GO:0042271 | susceptibility to natural killer cell mediated cytotoxicity | 5.97E-23 | Exhausted T cells |
| GO:0045879 | negative regulation of smoothened signaling pathway | 9.48E-22 | Effector T cells |
| GO:0051251 | positive regulation of lymphocyte activation | 1.42E-21 | Exhausted T cells |
| GO:0009756 | carbohydrate mediated signaling | 1.42E-21 | Exhausted T cells |
| GO:0016046 | detection of fungus | 8.52E-21 | Exhausted T cells |
| GO:0006509 | membrane protein ectodomain proteolysis | 1.65E-20 | Exhausted T cells |
| GO:0030449 | regulation of complement activation | 2.29E-20 | Exhausted T cells |
| GO:0080111 | DNA demethylation | 7.68E-20 | Exhausted T cells |
| GO:0045919 | positive regulation of cytolysis | 1.59E-19 | Effector T cells |
| GO:0045082 | positive regulation of interleukin-10 biosynthetic process | 1.02E-18 | Exhausted T cells |
| GO:0051563 | smooth endoplasmic reticulum calcium ion homeostasis | 3.89E-18 | Exhausted T cells |
| GO:0060267 | positive regulation of respiratory burst | 2.32E-17 | Exhausted T cells |
| GO:0051712 | positive regulation of killing of cells of another organism | 4.41E-17 | Exhausted T cells |
| GO:0021801 | cerebral cortex radial glia guided migration | 8.66E-17 | Effector T cells |
| GO:0019370 | leukotriene biosynthetic process | 1.06E-16 | Exhausted T cells |
| GO:0006910 | phagocytosis, recognition | 2.42E-16 | Exhausted T cells |
| GO:0032402 | melanosome transport | 2.53E-16 | Exhausted T cells |
| GO:0010573 | vascular endothelial growth factor production | 4.75E-16 | Effector T cells |
| GO:0042832 | defense response to protozoan | 1.52E-15 | Exhausted T cells |
| GO:0045084 | positive regulation of interleukin-12 biosynthetic process | 2.43E-15 | Exhausted T cells |
| GO:0051606 | detection of stimulus | 2.69E-15 | Exhausted T cells |
| GO:0030705 | cytoskeleton-dependent intracellular transport | 2.83E-15 | Exhausted T cells |
| GO:0040015 | negative regulation of multicellular organism growth | 4.13E-15 | Effector T cells |
| GO:0002221 | pattern recognition receptor signaling pathway | 4.11E-14 | Exhausted T cells |
| GO:0045416 | positive regulation of interleukin-8 biosynthetic process | 1.30E-13 | Exhausted T cells |
| GO:0045921 | positive regulation of exocytosis | 2.77E-13 | Exhausted T cells |
| GO:0006487 | protein N-linked glycosylation | 4.82E-13 | Exhausted T cells |
| GO:0035567 | non-canonical Wnt receptor signaling pathway | 7.94E-13 | Exhausted T cells |
| GO:0070528 | protein kinase C signaling cascade | 1.16E-12 | Effector T cells |
| GO:0046368 | GDP-L-fucose metabolic process | 2.86E-12 | Exhausted T cells |
| GO:0006935 | chemotaxis | 6.79E-12 | Effector T cells |
| GO:0032731 | positive regulation of interleukin-1 beta production | 6.85E-12 | Exhausted T cells |
| GO:0046370 | fructose biosynthetic process | 7.13E-12 | Exhausted T cells |
| GO:0032945 | negative regulation of mononuclear cell proliferation | 1.05E-11 | Effector T cells |
| GO:0033578 | protein glycosylation in Golgi | 1.43E-11 | Exhausted T cells |
| GO:0032692 | negative regulation of interleukin-1 production | 2.19E-11 | Exhausted T cells |
| GO:0032703 | negative regulation of interleukin-2 production | 3.69E-11 | Exhausted T cells |
| GO:0090303 | positive regulation of wound healing | 4.16E-11 | Exhausted T cells |
| GO:0032400 | melanosome localization | 4.87E-11 | Exhausted T cells |
| GO:0002725 | negative regulation of T cell cytokine production | 5.06E-11 | Exhausted T cells |
| GO:0046636 | negative regulation of alpha-beta T cell activation | 6.26E-11 | Effector T cells |
| GO:0060312 | regulation of blood vessel remodeling | 6.55E-11 | Exhausted T cells |
| GO:0045601 | regulation of endothelial cell differentiation | 6.55E-11 | Exhausted T cells |
| GO:0010759 | positive regulation of macrophage chemotaxis | 9.23E-11 | Effector T cells |
| GO:0032831 | positive regulation of CD4-positive, CD25-positive, alpha-beta regulatory T cell differentiation | 1.36E-10 | Exhausted T cells |
| GO:0050435 | beta-amyloid metabolic process | 1.46E-10 | Exhausted T cells |
| GO:0045410 | positive regulation of interleukin-6 biosynthetic process | 2.15E-10 | Exhausted T cells |
| GO:0016339 | calcium-dependent cell-cell adhesion | 2.32E-10 | Exhausted T cells |
| GO:0042534 | regulation of tumor necrosis factor biosynthetic process | 2.75E-10 | Effector T cells |
| GO:0035025 | positive regulation of Rho protein signal transduction | 4.73E-10 | Effector T cells |
| GO:0060828 | regulation of canonical Wnt receptor signaling pathway | 5.99E-10 | Exhausted T cells |
| GO:0006977 | DNA damage response, signal transduction by p53 class mediator resulting in cell cycle arrest | 9.35E-10 | Exhausted T cells |
| GO:0006887 | exocytosis | 9.65E-10 | Exhausted T cells |
| GO:0030195 | negative regulation of blood coagulation | 1.26E-09 | Exhausted T cells |
| GO:0014063 | negative regulation of serotonin secretion | 1.31E-09 | Effector T cells |
| GO:0051055 | negative regulation of lipid biosynthetic process | 1.32E-09 | Exhausted T cells |
| GO:0006491 | N-glycan processing | 1.37E-09 | Exhausted T cells |
| GO:0032689 | negative regulation of interferon-gamma production | 1.60E-09 | Common |
| GO:0070498 | interleukin-1-mediated signaling pathway | 1.81E-09 | Exhausted T cells |
| GO:0045077 | negative regulation of interferon-gamma biosynthetic process | 2.61E-09 | Effector T cells |
| GO:0045636 | positive regulation of melanocyte differentiation | 2.74E-09 | Effector T cells |
| GO:0002667 | regulation of T cell anergy | 4.27E-09 | Exhausted T cells |
| GO:0034699 | response to luteinizing hormone stimulus | 5.74E-09 | Effector T cells |
| GO:0002309 | T cell proliferation involved in immune response | 8.16E-09 | Effector T cells |
| GO:0042982 | amyloid precursor protein metabolic process | 8.90E-09 | Exhausted T cells |
| GO:0010875 | positive regulation of cholesterol efflux | 9.01E-09 | Effector T cells |
| GO:0035108 | limb morphogenesis | 9.01E-09 | Effector T cells |
| GO:0042307 | positive regulation of protein import into nucleus | 9.67E-09 | Exhausted T cells |
| GO:0030853 | negative regulation of granulocyte differentiation | 1.40E-08 | Exhausted T cells |
| GO:0015871 | choline transport | 1.46E-08 | Exhausted T cells |
| GO:0006641 | triglyceride metabolic process | 1.56E-08 | Exhausted T cells |
| GO:0042355 | L-fucose catabolic process | 1.70E-08 | Exhausted T cells |
| GO:0008589 | regulation of smoothened signaling pathway | 1.78E-08 | Effector T cells |
| GO:0070588 | calcium ion transmembrane transport | 1.81E-08 | Exhausted T cells |
| GO:0030326 | embryonic limb morphogenesis | 1.84E-08 | Effector T cells |
| GO:0030099 | myeloid cell differentiation | 1.99E-08 | Exhausted T cells |
| GO:0031293 | membrane protein intracellular domain proteolysis | 2.15E-08 | Exhausted T cells |
| GO:0043542 | endothelial cell migration | 2.29E-08 | Effector T cells |
| GO:0060086 | circadian temperature homeostasis | 2.29E-08 | Exhausted T cells |
| GO:0060318 | definitive erythrocyte differentiation | 2.86E-08 | Effector T cells |
| GO:0045209 | MAPK phosphatase export from nucleus, leptomycin B sensitive | 3.42E-08 | Exhausted T cells |
| GO:0008037 | cell recognition | 3.47E-08 | Exhausted T cells |
| GO:0090311 | regulation of protein deacetylation | 3.63E-08 | Exhausted T cells |
| GO:0031532 | actin cytoskeleton reorganization | 3.66E-08 | Exhausted T cells |
| GO:0031161 | phosphatidylinositol catabolic process | 5.66E-08 | Exhausted T cells |
| GO:0015728 | mevalonate transport | 6.24E-08 | Exhausted T cells |
| GO:0010594 | regulation of endothelial cell migration | 7.46E-08 | Exhausted T cells |
| GO:0042325 | regulation of phosphorylation | 9.51E-08 | Exhausted T cells |
| GO:0008544 | epidermis development | 1.10E-07 | Exhausted T cells |
| GO:0002740 | negative regulation of cytokine secretion involved in immune response | 1.18E-07 | Effector T cells |
| GO:0007018 | microtubule-based movement | 1.27E-07 | Exhausted T cells |
| GO:0045085 | negative regulation of interleukin-2 biosynthetic process | 1.70E-07 | Exhausted T cells |
| GO:0042536 | negative regulation of tumor necrosis factor biosynthetic process | 2.05E-07 | Effector T cells |
| GO:0032609 | interferon-gamma production | 2.05E-07 | Effector T cells |
| GO:0045892 | negative regulation of transcription, DNA-dependent | 2.17E-07 | Exhausted T cells |
| GO:0048261 | negative regulation of receptor-mediated endocytosis | 2.27E-07 | Exhausted T cells |
| GO:0016188 | synaptic vesicle maturation | 2.37E-07 | Exhausted T cells |
| GO:0045821 | positive regulation of glycolysis | 2.40E-07 | Exhausted T cells |
| GO:0007270 | nerve-nerve synaptic transmission | 2.54E-07 | Exhausted T cells |
| GO:0045428 | regulation of nitric oxide biosynthetic process | 2.57E-07 | Effector T cells |
| GO:0045785 | positive regulation of cell adhesion | 2.58E-07 | Effector T cells |
| GO:0031334 | positive regulation of protein complex assembly | 3.35E-07 | Exhausted T cells |
| GO:0043116 | negative regulation of vascular permeability | 3.43E-07 | Exhausted T cells |
| GO:0043280 | positive regulation of caspase activity | 3.67E-07 | Exhausted T cells |
| GO:0032354 | response to follicle-stimulating hormone stimulus | 3.98E-07 | Effector T cells |
| GO:0030317 | sperm motility | 4.20E-07 | Exhausted T cells |
| GO:0043306 | positive regulation of mast cell degranulation | 4.52E-07 | Effector T cells |
| GO:0042327 | positive regulation of phosphorylation | 4.55E-07 | Exhausted T cells |
| GO:0047484 | regulation of response to osmotic stress | 4.87E-07 | Effector T cells |
| GO:0045204 | MAPK export from nucleus | 5.09E-07 | Exhausted T cells |
| GO:0006637 | acyl-CoA metabolic process | 5.59E-07 | Exhausted T cells |
| GO:0006879 | cellular iron ion homeostasis | 6.27E-07 | Exhausted T cells |
| GO:0010757 | negative regulation of plasminogen activation | 6.63E-07 | Exhausted T cells |
| GO:0032713 | negative regulation of interleukin-4 production | 8.05E-07 | Exhausted T cells |
| GO:0071526 | semaphorin-plexin signaling pathway | 8.83E-07 | Exhausted T cells |
| GO:0045429 | positive regulation of nitric oxide biosynthetic process | 9.99E-07 | Exhausted T cells |
| GO:0016055 | Wnt receptor signaling pathway | 1.09E-06 | Exhausted T cells |
| GO:0034695 | response to prostaglandin E stimulus | 1.19E-06 | Effector T cells |
| GO:0045078 | positive regulation of interferon-gamma biosynthetic process | 1.20E-06 | Exhausted T cells |
| GO:0050868 | negative regulation of T cell activation | 1.24E-06 | Exhausted T cells |
| GO:0042036 | negative regulation of cytokine biosynthetic process | 1.24E-06 | Exhausted T cells |
| GO:0060544 | regulation of necroptosis | 1.26E-06 | Exhausted T cells |
| GO:0048143 | astrocyte activation | 1.29E-06 | Exhausted T cells |
| GO:0045668 | negative regulation of osteoblast differentiation | 1.54E-06 | Effector T cells |
| GO:0007050 | cell cycle arrest | 1.61E-06 | Exhausted T cells |
| GO:0042058 | regulation of epidermal growth factor receptor signaling pathway | 1.63E-06 | Exhausted T cells |
| GO:0043085 | positive regulation of catalytic activity | 1.66E-06 | Exhausted T cells |
| GO:0006631 | fatty acid metabolic process | 1.95E-06 | Exhausted T cells |
| GO:0001915 | negative regulation of T cell mediated cytotoxicity | 2.66E-06 | Common |
| GO:0060071 | Wnt receptor signaling pathway, planar cell polarity pathway | 3.09E-06 | Exhausted T cells |
| GO:0045944 | positive regulation of transcription from RNA polymerase II promoter | 3.37E-06 | Exhausted T cells |
| GO:0048489 | synaptic vesicle transport | 3.40E-06 | Exhausted T cells |
| GO:0009312 | oligosaccharide biosynthetic process | 3.96E-06 | Exhausted T cells |
| GO:0045453 | bone resorption | 4.22E-06 | Exhausted T cells |
| GO:0015914 | phospholipid transport | 4.35E-06 | Effector T cells |
| GO:0032691 | negative regulation of interleukin-1 beta production | 4.59E-06 | Effector T cells |
| GO:0050853 | B cell receptor signaling pathway | 4.66E-06 | Exhausted T cells |
| GO:0046466 | membrane lipid catabolic process | 4.80E-06 | Exhausted T cells |
| GO:0045717 | negative regulation of fatty acid biosynthetic process | 4.81E-06 | Exhausted T cells |
| GO:0002517 | T cell tolerance induction | 4.95E-06 | Exhausted T cells |
| GO:0032700 | negative regulation of interleukin-17 production | 5.39E-06 | Exhausted T cells |
| GO:0060907 | positive regulation of macrophage cytokine production | 6.48E-06 | Effector T cells |
| GO:0050764 | regulation of phagocytosis | 6.52E-06 | Effector T cells |
| GO:0051926 | negative regulation of calcium ion transport | 7.23E-06 | Effector T cells |
| GO:0007596 | blood coagulation | 7.69E-06 | Exhausted T cells |
| GO:0043517 | positive regulation of DNA damage response, signal transduction by p53 class mediator | 8.06E-06 | Exhausted T cells |
| GO:0045806 | negative regulation of endocytosis | 8.19E-06 | Effector T cells |
| GO:0090331 | negative regulation of platelet aggregation | 1.05E-05 | Exhausted T cells |
| GO:0045954 | positive regulation of natural killer cell mediated cytotoxicity | 1.09E-05 | Exhausted T cells |
| GO:0031064 | negative regulation of histone deacetylation | 1.16E-05 | Exhausted T cells |
| GO:0045556 | positive regulation of TRAIL biosynthetic process | 1.16E-05 | Exhausted T cells |
| GO:0050860 | negative regulation of T cell receptor signaling pathway | 1.20E-05 | Exhausted T cells |
| GO:0050776 | regulation of immune response | 1.46E-05 | Common |
| GO:0002768 | immune response-regulating cell surface receptor signaling pathway | 1.71E-05 | Effector T cells |
| GO:0010611 | regulation of cardiac muscle hypertrophy | 1.86E-05 | Exhausted T cells |
| GO:0046683 | response to organophosphorus | 1.99E-05 | Exhausted T cells |
| GO:0043331 | response to dsRNA | 2.30E-05 | Exhausted T cells |
| GO:0007167 | enzyme linked receptor protein signaling pathway | 2.43E-05 | Effector T cells |
| GO:0007267 | cell-cell signaling | 2.54E-05 | Effector T cells |
| GO:0018279 | protein N-linked glycosylation via asparagine | 2.64E-05 | Exhausted T cells |
| GO:0002767 | immune response-inhibiting cell surface receptor signaling pathway | 2.69E-05 | Effector T cells |
| GO:0001676 | long-chain fatty acid metabolic process | 2.81E-05 | Exhausted T cells |
| GO:0050777 | negative regulation of immune response | 3.19E-05 | Exhausted T cells |
| GO:0051271 | negative regulation of cellular component movement | 3.29E-05 | Effector T cells |
| GO:0014808 | release of sequestered calcium ion into cytosol by sarcoplasmic reticulum | 3.51E-05 | Exhausted T cells |
| GO:0031398 | positive regulation of protein ubiquitination | 3.60E-05 | Exhausted T cells |
| GO:0006911 | phagocytosis, engulfment | 3.63E-05 | Effector T cells |
| GO:0060347 | heart trabecula formation | 3.92E-05 | Effector T cells |
| GO:0050808 | synapse organization | 6.04E-05 | Exhausted T cells |
| GO:0045086 | positive regulation of interleukin-2 biosynthetic process | 6.06E-05 | Exhausted T cells |
| GO:0050715 | positive regulation of cytokine secretion | 6.20E-05 | Effector T cells |
| GO:0014066 | regulation of phosphoinositide 3-kinase cascade | 6.23E-05 | Exhausted T cells |
| GO:0050901 | leukocyte tethering or rolling | 6.79E-05 | Effector T cells |
| GO:0060352 | cell adhesion molecule production | 7.11E-05 | Exhausted T cells |
| GO:0045590 | negative regulation of regulatory T cell differentiation | 7.24E-05 | Exhausted T cells |
| GO:0043271 | negative regulation of ion transport | 7.29E-05 | Exhausted T cells |
| GO:0070098 | chemokine-mediated signaling pathway | 7.42E-05 | Effector T cells |
| GO:0019563 | glycerol catabolic process | 7.89E-05 | Exhausted T cells |
| GO:0002430 | complement receptor mediated signaling pathway | 8.29E-05 | Effector T cells |
| GO:0007200 | activation of phospholipase C activity by G-protein coupled receptor protein signaling pathway coupled to IP3 second messenger | 8.32E-05 | Effector T cells |
| GO:0008089 | anterograde axon cargo transport | 9.03E-05 | Exhausted T cells |
| GO:0007266 | Rho protein signal transduction | 9.06E-05 | Effector T cells |
| GO:0000188 | inactivation of MAPK activity | 9.07E-05 | Exhausted T cells |
| GO:0006897 | endocytosis | 9.59E-05 | Exhausted T cells |
