## Supplementary material for "Single-cell Long Non-coding RNA Landscape of T Cells in Human Cancer Immunity": Table S11.docx

**Table S11 Functional enrichment results of CD4 Treg signature lncRNAs**

| **GO ID** | **GO name** | **Adjusted *P*-value** |
| --- | --- | --- |
| GO:0007220 | Notch receptor processing | 1.79E-26 |
| GO:0023052 | signaling | 3.19E-26 |
| GO:0032468 | Golgi calcium ion homeostasis | 2.80E-25 |
| GO:0008154 | actin polymerization or depolymerization | 1.71E-24 |
| GO:0032472 | Golgi calcium ion transport | 3.35E-24 |
| GO:0042271 | susceptibility to natural killer cell mediated cytotoxicity | 7.96E-23 |
| GO:0032491 | detection of molecule of fungal origin | 1.65E-22 |
| GO:0009756 | carbohydrate mediated signaling | 1.81E-21 |
| GO:0051251 | positive regulation of lymphocyte activation | 1.81E-21 |
| GO:0016046 | detection of fungus | 1.08E-20 |
| GO:0080111 | DNA demethylation | 2.81E-20 |
| GO:0051209 | release of sequestered calcium ion into cytosol | 1.65E-19 |
| GO:0045082 | positive regulation of interleukin-10 biosynthetic process | 1.29E-18 |
| GO:0051563 | smooth endoplasmic reticulum calcium ion homeostasis | 1.94E-18 |
| GO:0071421 | manganese ion transmembrane transport | 3.46E-18 |
| GO:0060267 | positive regulation of respiratory burst | 2.94E-17 |
| GO:0071226 | cellular response to molecule of fungal origin | 2.94E-17 |
| GO:0051712 | positive regulation of killing of cells of other organism | 5.60E-17 |
| GO:0019370 | leukotriene biosynthetic process | 8.47E-17 |
| GO:0006910 | phagocytosis, recognition | 3.08E-16 |
| GO:0071639 | positive regulation of monocyte chemotactic protein-1 production | 5.12E-16 |
| GO:0045084 | positive regulation of interleukin-12 biosynthetic process | 3.09E-15 |
| GO:0051606 | detection of stimulus | 3.57E-15 |
| GO:0042832 | defense response to protozoan | 2.59E-14 |
| GO:0045078 | positive regulation of interferon-gamma biosynthetic process | 3.49E-14 |
| GO:0002221 | pattern recognition receptor signaling pathway | 5.22E-14 |
| GO:0000435 | positive regulation of transcription from RNA polymerase II promoter by galactose | 6.12E-14 |
| GO:0045086 | positive regulation of interleukin-2 biosynthetic process | 8.63E-14 |
| GO:0006509 | membrane protein ectodomain proteolysis | 1.46E-13 |
| GO:0045416 | positive regulation of interleukin-8 biosynthetic process | 1.66E-13 |
| GO:0038016 | insulin receptor internalization | 1.29E-12 |
| GO:0002265 | astrocyte activation involved in immune response | 1.51E-12 |
| GO:2000582 | positive regulation of plus-end-directed microtubule motor activity | 2.07E-12 |
| GO:1900017 | positive regulation of cytokine production involved in inflammatory response | 2.25E-12 |
| GO:0035333 | Notch receptor processing, ligand-dependent | 2.44E-12 |
| GO:0005977 | glycogen metabolic process | 2.87E-12 |
| GO:0032731 | positive regulation of interleukin-1 beta production | 8.71E-12 |
| GO:0031529 | ruffle organization | 8.85E-12 |
| GO:0002859 | negative regulation of natural killer cell mediated cytotoxicity directed against tumor cell target | 9.01E-12 |
| GO:0032703 | negative regulation of interleukin-2 production | 3.38E-11 |
| GO:0032692 | negative regulation of interleukin-1 production | 3.59E-11 |
| GO:0038158 | granulocyte colony-stimulating factor signaling pathway | 3.59E-11 |
| GO:1901143 | insulin catabolic process | 3.59E-11 |
| GO:0002725 | negative regulation of T cell cytokine production | 4.64E-11 |
| GO:0090303 | positive regulation of wound healing | 5.29E-11 |
| GO:0050435 | beta-amyloid metabolic process | 7.29E-11 |
| GO:0060312 | regulation of blood vessel remodeling | 1.07E-10 |
| GO:0043318 | negative regulation of cytotoxic T cell degranulation | 1.07E-10 |
| GO:0045601 | regulation of endothelial cell differentiation | 1.07E-10 |
| GO:0032930 | positive regulation of superoxide anion generation | 1.24E-10 |
| GO:0032831 | positive regulation of CD4-positive, CD25-positive, alpha-beta regulatory T cell differentiation | 1.25E-10 |
| GO:0016339 | calcium-dependent cell-cell adhesion | 1.39E-10 |
| GO:0045410 | positive regulation of interleukin-6 biosynthetic process | 2.73E-10 |
| GO:0060828 | regulation of canonical Wnt signaling pathway | 3.00E-10 |
| GO:0006977 | DNA damage response, signal transduction by p53 class mediator resulting in cell cycle arrest | 1.17E-09 |
| GO:0030195 | negative regulation of blood coagulation | 1.30E-09 |
| GO:0071313 | cellular response to caffeine | 1.38E-09 |
| GO:0032689 | negative regulation of interferon-gamma production | 1.47E-09 |
| GO:0051055 | negative regulation of lipid biosynthetic process | 2.16E-09 |
| GO:2000346 | negative regulation of hepatocyte proliferation | 2.16E-09 |
| GO:0070498 | interleukin-1-mediated signaling pathway | 2.26E-09 |
| GO:0002667 | regulation of T cell anergy | 3.91E-09 |
| GO:0042982 | amyloid precursor protein metabolic process | 4.45E-09 |
| GO:0042307 | positive regulation of protein import into nucleus | 4.86E-09 |
| GO:0050766 | positive regulation of phagocytosis | 6.46E-09 |
| GO:0097112 | gamma-aminobutyric acid receptor clustering | 6.58E-09 |
| GO:0030099 | myeloid cell differentiation | 7.59E-09 |
| GO:0035726 | common myeloid progenitor cell proliferation | 9.96E-09 |
| GO:0031293 | membrane protein intracellular domain proteolysis | 1.07E-08 |
| GO:0006816 | calcium ion transport | 1.35E-08 |
| GO:0031532 | actin cytoskeleton reorganization | 2.20E-08 |
| GO:0030853 | negative regulation of granulocyte differentiation | 2.29E-08 |
| GO:0070588 | calcium ion transmembrane transport | 2.98E-08 |
| GO:0000188 | inactivation of MAPK activity | 3.19E-08 |
| GO:0002362 | CD4-positive, CD25-positive, alpha-beta regulatory T cell lineage commitment | 3.30E-08 |
| GO:0097494 | regulation of vesicle size | 3.30E-08 |
| GO:0060086 | circadian temperature homeostasis | 4.46E-08 |
| GO:0008037 | cell recognition | 4.61E-08 |
| GO:0031161 | phosphatidylinositol catabolic process | 4.66E-08 |
| GO:0042325 | regulation of phosphorylation | 4.75E-08 |
| GO:0008544 | epidermis development | 6.62E-08 |
| GO:0031334 | positive regulation of protein complex assembly | 8.40E-08 |
| GO:0042130 | negative regulation of T cell proliferation | 8.73E-08 |
| GO:2001214 | positive regulation of vasculogenesis | 9.06E-08 |
| GO:0007018 | microtubule-based movement | 9.79E-08 |
| GO:0045821 | positive regulation of glycolysis | 1.20E-07 |
| GO:0010594 | regulation of endothelial cell migration | 1.22E-07 |
| GO:0030299 | intestinal cholesterol absorption | 1.24E-07 |
| GO:0045085 | negative regulation of interleukin-2 biosynthetic process | 1.56E-07 |
| GO:0071812 | positive regulation of fever generation by positive regulation of prostaglandin secretion | 1.62E-07 |
| GO:0045892 | negative regulation of transcription, DNA-templated | 1.63E-07 |
| GO:0048261 | negative regulation of receptor-mediated endocytosis | 1.67E-07 |
| GO:0016188 | synaptic vesicle maturation | 1.70E-07 |
| GO:0042327 | positive regulation of phosphorylation | 2.27E-07 |
| GO:0043280 | positive regulation of cysteine-type endopeptidase activity involved in apoptotic process | 4.68E-07 |
| GO:0045860 | positive regulation of protein kinase activity | 4.99E-07 |
| GO:0043116 | negative regulation of vascular permeability | 5.62E-07 |
| GO:0048143 | astrocyte activation | 6.46E-07 |
| GO:0010757 | negative regulation of plasminogen activation | 7.02E-07 |
| GO:0035459 | cargo loading into vesicle | 7.14E-07 |
| GO:0032713 | negative regulation of interleukin-4 production | 7.38E-07 |
| GO:0006879 | cellular iron ion homeostasis | 1.04E-06 |
| GO:0042036 | negative regulation of cytokine biosynthetic process | 1.14E-06 |
| GO:0050868 | negative regulation of T cell activation | 1.24E-06 |
| GO:0045429 | positive regulation of nitric oxide biosynthetic process | 1.27E-06 |
| GO:0006631 | fatty acid metabolic process | 1.95E-06 |
| GO:0007050 | cell cycle arrest | 2.41E-06 |
| GO:0042058 | regulation of epidermal growth factor receptor signaling pathway | 2.66E-06 |
| GO:0048013 | ephrin receptor signaling pathway | 3.84E-06 |
| GO:0046466 | membrane lipid catabolic process | 3.84E-06 |
| GO:0001915 | negative regulation of T cell mediated cytotoxicity | 4.35E-06 |
| GO:1901895 | negative regulation of calcium-transporting ATPase activity | 4.53E-06 |
| GO:0032700 | negative regulation of interleukin-17 production | 4.94E-06 |
| GO:0045717 | negative regulation of fatty acid biosynthetic process | 7.87E-06 |
| GO:1901315 | negative regulation of histone H2A K63-linked ubiquitination | 9.42E-06 |
| GO:0007040 | lysosome organization | 1.03E-05 |
| GO:0031064 | negative regulation of histone deacetylation | 1.06E-05 |
| GO:0045954 | positive regulation of natural killer cell mediated cytotoxicity | 1.09E-05 |
| GO:0071848 | positive regulation of ERK1 and ERK2 cascade via TNFSF11-mediated signaling | 1.10E-05 |
| GO:1902004 | positive regulation of beta-amyloid formation | 1.32E-05 |
| GO:0097213 | regulation of lysosomal membrane permeability | 1.67E-05 |
| GO:0010611 | regulation of cardiac muscle hypertrophy | 1.86E-05 |
| GO:0046683 | response to organophosphorus | 1.93E-05 |
| GO:0050860 | negative regulation of T cell receptor signaling pathway | 1.95E-05 |
| GO:0090331 | negative regulation of platelet aggregation | 2.06E-05 |
| GO:0016185 | synaptic vesicle budding from presynaptic membrane | 2.43E-05 |
| GO:1990108 | protein linear deubiquitination | 2.46E-05 |
| GO:0006486 | protein glycosylation | 2.59E-05 |
| GO:0014808 | release of sequestered calcium ion into cytosol by sarcoplasmic reticulum | 2.66E-05 |
| GO:0001676 | long-chain fatty acid metabolic process | 2.93E-05 |
| GO:0050808 | synapse organization | 3.02E-05 |
| GO:2000009 | negative regulation of protein localization to cell surface | 3.14E-05 |
| GO:0030334 | regulation of cell migration | 3.93E-05 |
| GO:0071847 | TNFSF11-mediated signaling pathway | 4.28E-05 |
| GO:0007010 | cytoskeleton organization | 5.37E-05 |
| GO:0043433 | negative regulation of sequence-specific DNA binding transcription factor activity | 5.55E-05 |
| GO:0006937 | regulation of muscle contraction | 6.11E-05 |
| GO:0032092 | positive regulation of protein binding | 6.23E-05 |
| GO:0044255 | cellular lipid metabolic process | 7.01E-05 |
| GO:0060352 | cell adhesion molecule production | 7.11E-05 |
| GO:0006493 | protein O-linked glycosylation | 7.41E-05 |
| GO:0006805 | xenobiotic metabolic process | 7.50E-05 |
| GO:0006897 | endocytosis | 8.03E-05 |
| GO:0045893 | positive regulation of transcription, DNA-templated | 8.18E-05 |
| GO:0007155 | cell adhesion | 9.43E-05 |
