## Supplementary figures and images for "Single-cell Long Non-coding RNA Landscape of T Cells in Human Cancer Immunity"

### Figure S1.jpg

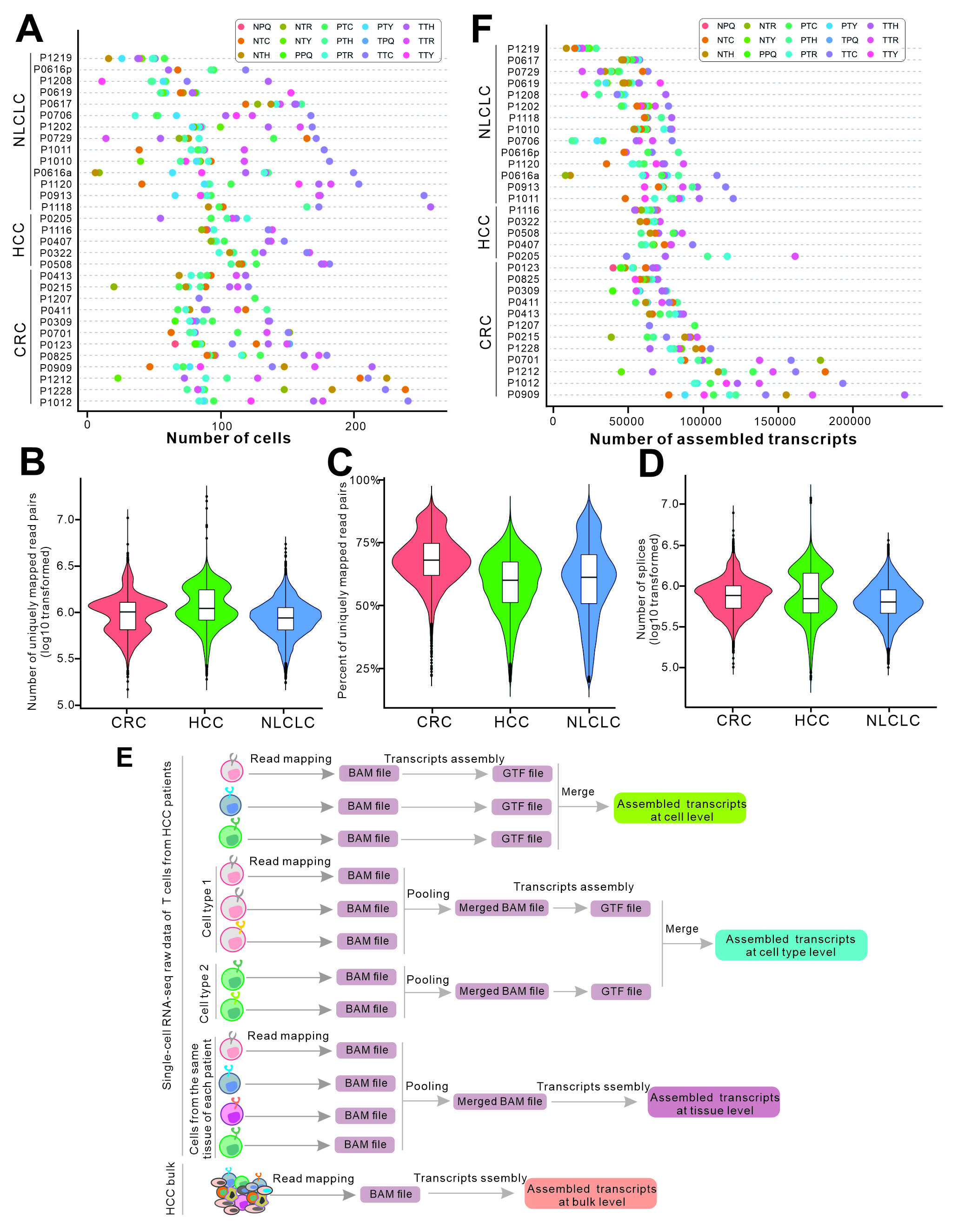

### Figure S2.jpg

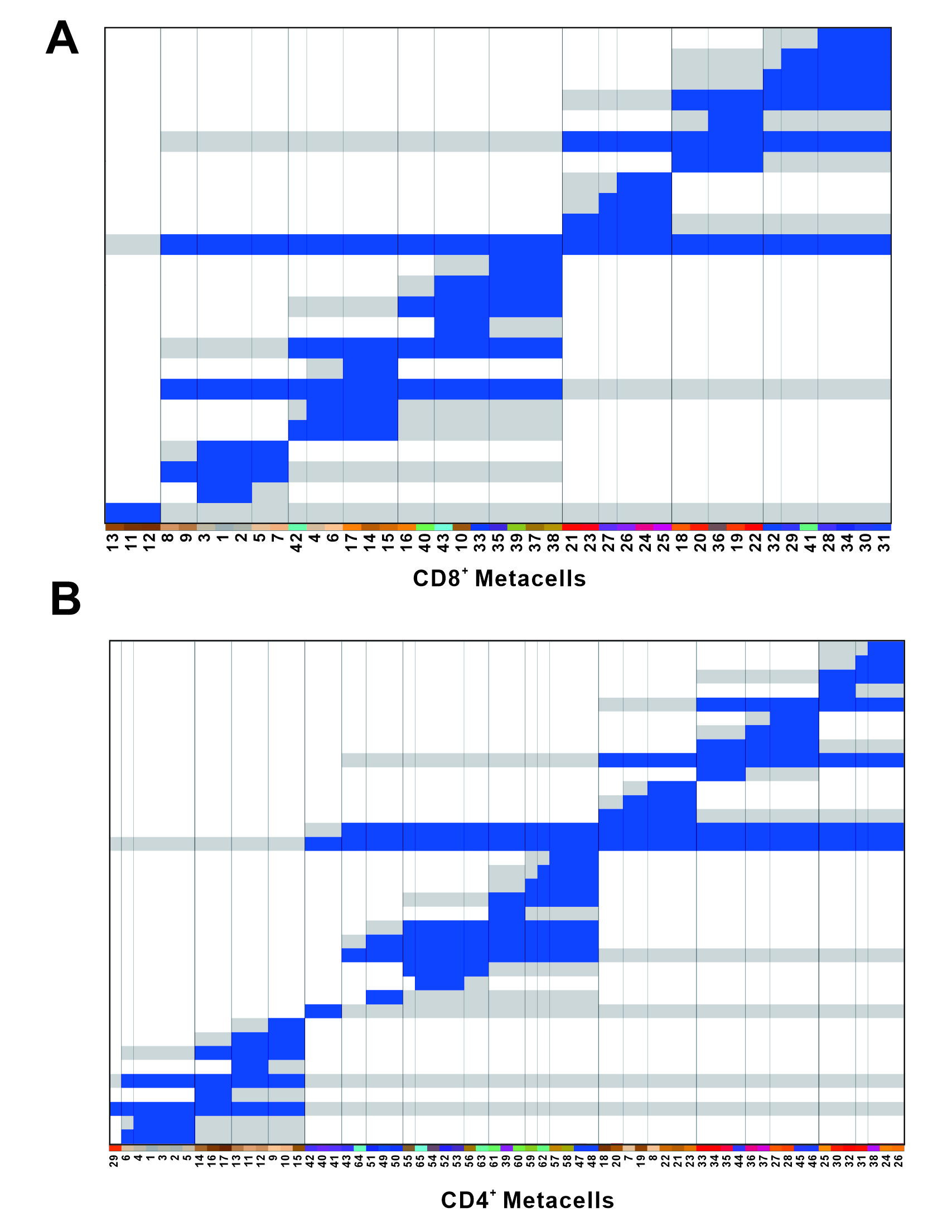

### Figure S3.jpg

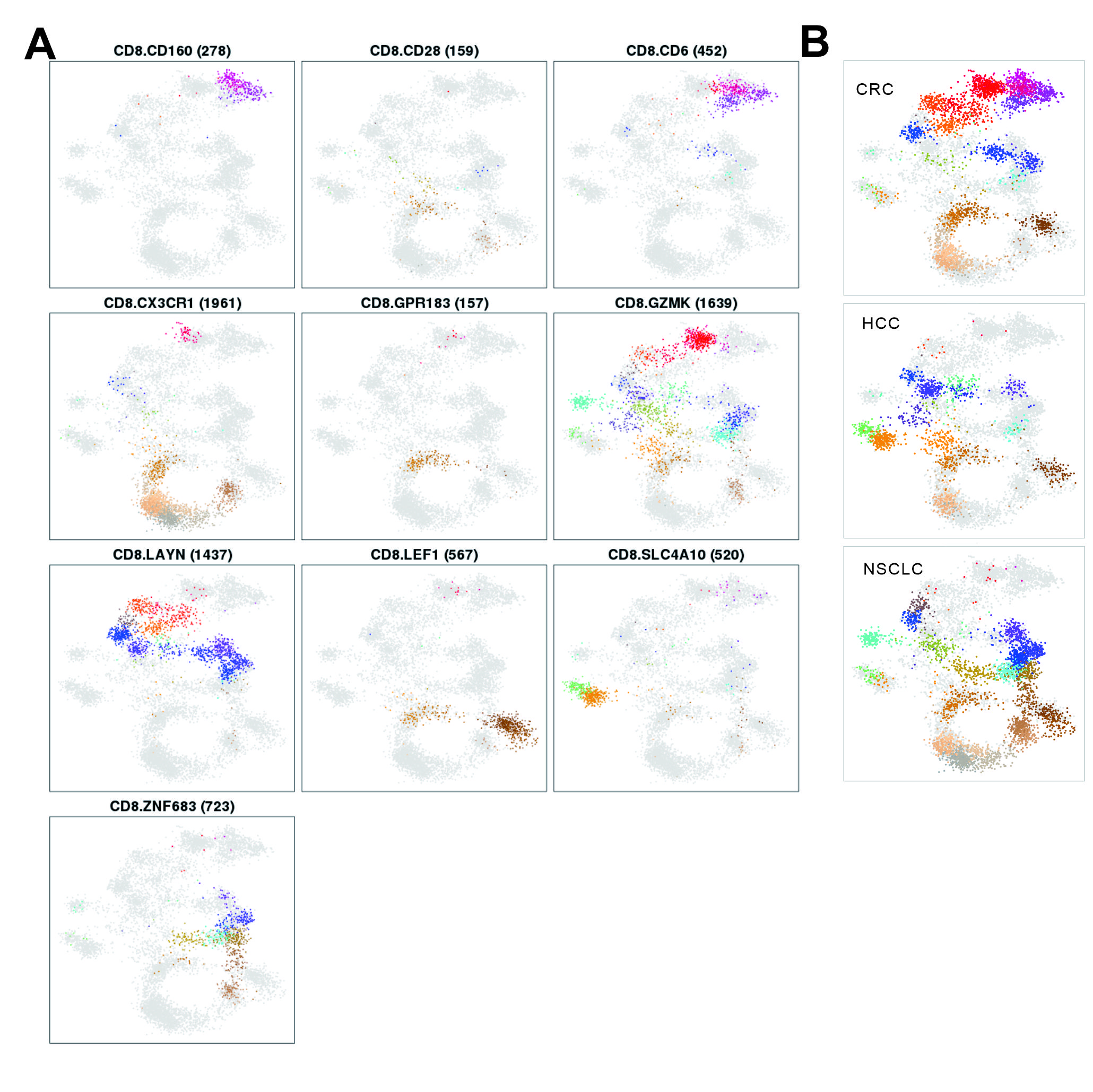

### Figure S4.jpg

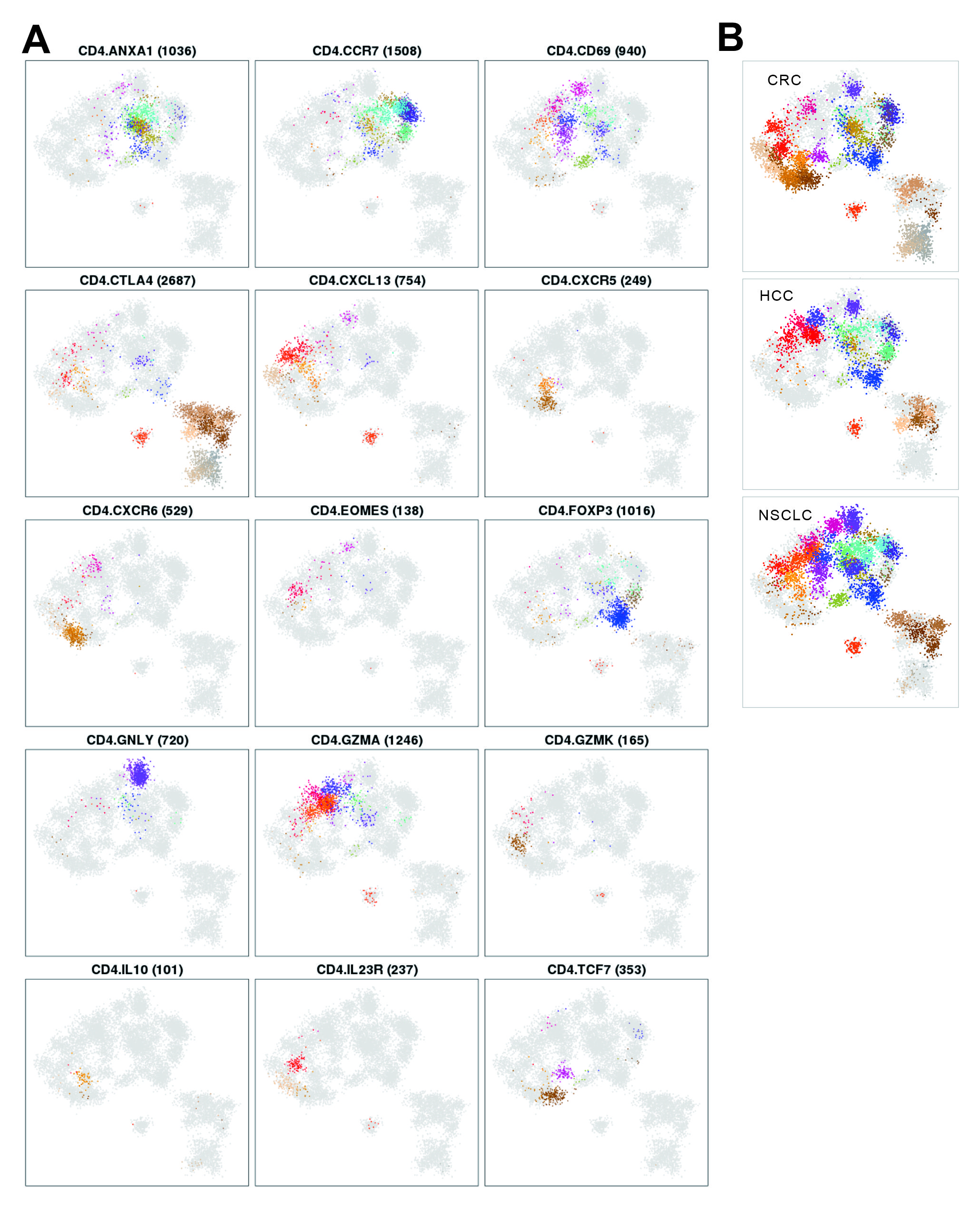

### Figure S5.jpg

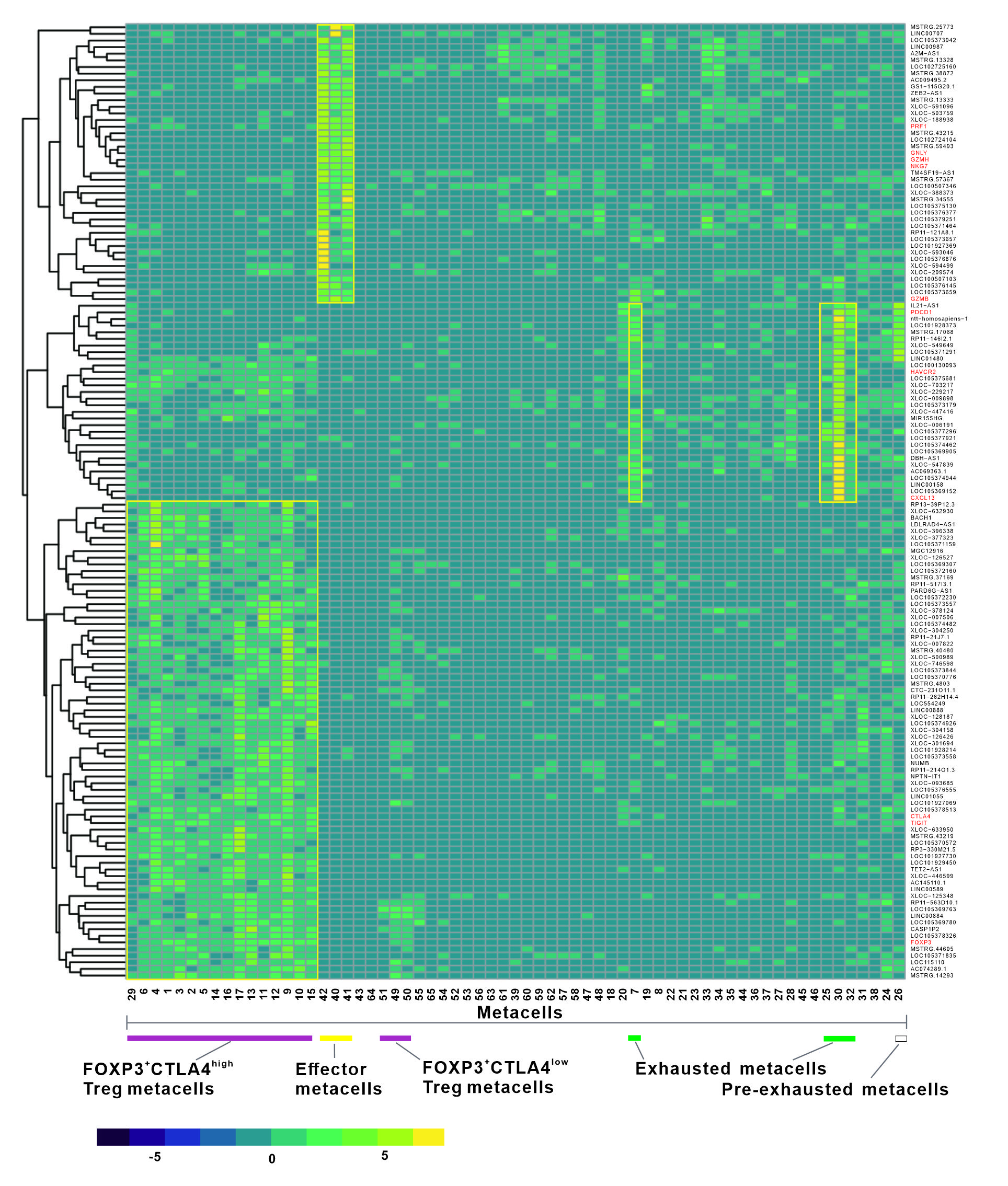

### Figure S6.jpg

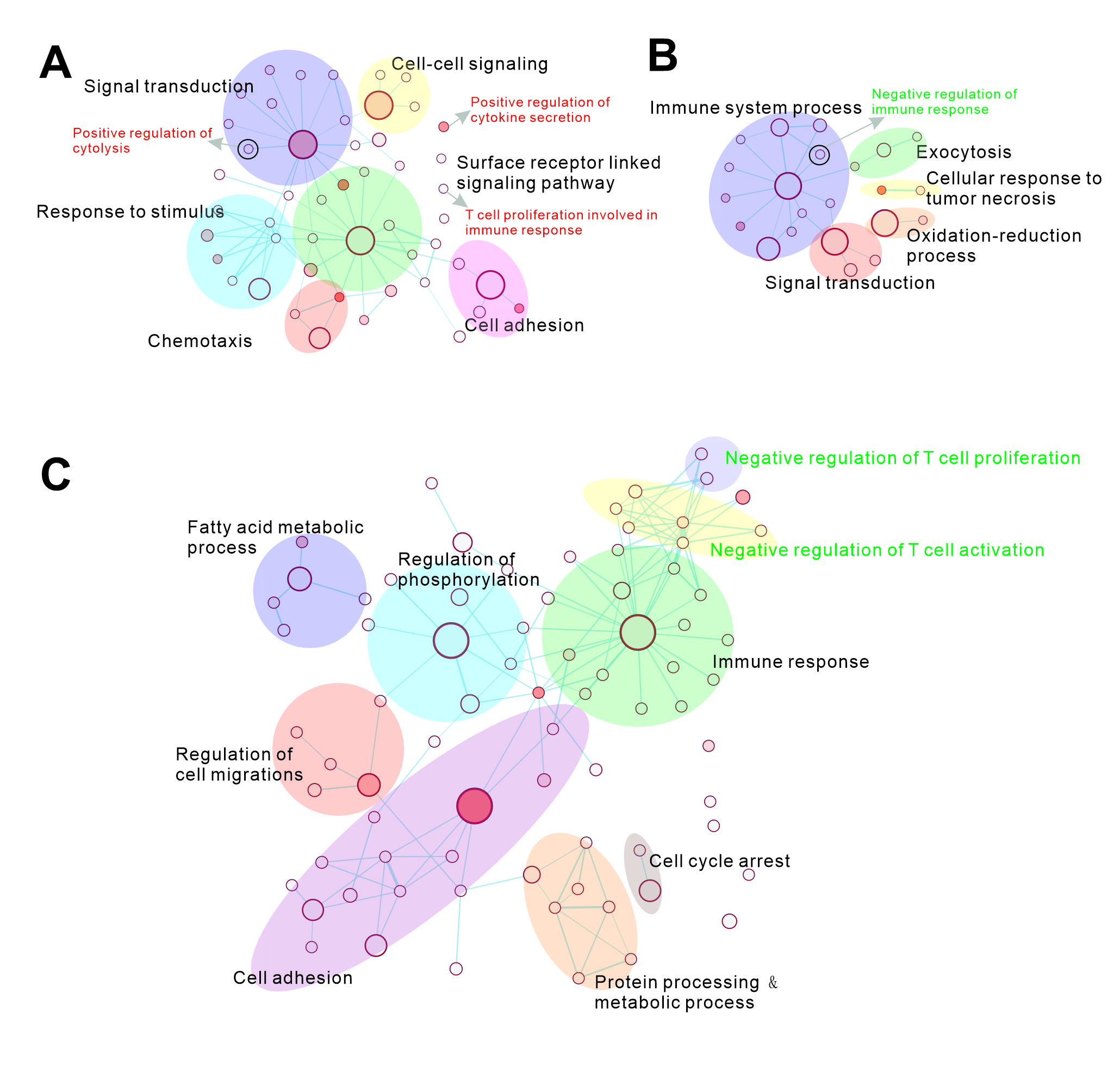
